# Hitchhiking on condensed chromatin promotes plasmid persistence in yeast without perturbing chromosome function

**DOI:** 10.1101/2020.06.08.139568

**Authors:** Hemant Kumar Prajapati, Deepanshu Kumar, Xian-Mei Yang, Chien-Hui Ma, Priyanka Mittal, Makkuni Jayaram, Santanu K. Ghosh

## Abstract

Equipartitioning by chromosome hitchhiking and copy number correction by DNA amplification are at the heart of the evolutionary success of the selfish yeast 2-micron plasmid. The present analysis reveals plasmid presence near centromeres and telomeres in mitotic cells, with a preference towards the latter. The observed correlation of plasmid missegregation with non-disjunction of rDNA and telomeres under Cdc14 inactivation, higher plasmid missegregation upon induced missegregation of chromosome XII but not chromosome III, requirement of condensin for plasmid stability and the interaction of the condensin subunit Brn1 with the plasmid partitioning system lend functional credence to condensed chromatin being favored for plasmid tethering. By homing to condensed/quiescent chromosome locales, and not over-perturbing genome homeostasis, the plasmid may minimize fitness conflicts with its host. Analogous persistence strategies may be utilized by other extrachromosomal selfish genomes, for example, episomes of mammalian viruses that also hitchhike on host chromosomes for their stable maintenance.

## Introduction

Eukaryotes, in contrast to prokaryotes, rarely harbor stably propagating extra-chromosomal DNA elements in their nuclei. However, circular plasmids have been identified in the budding yeast lineage and in the slime mold *Dictyostelium* (Broach, 1982; Hughes & Welker, 1999). In addition, papilloma and gammaherpes viruses exist as episomes in infected mammalian cells during long periods of latency (Coursey & McBride, 2019; Frappier, 2004; McBride, 2008). Eukaryotic nuclei also contain extra-chromosomal circular DNA molecules, called eccDNA that are excised from chromosomes (Cohen & Segal, 2009; Møller et al., 2015; Shibata et al., 2012). Their formation, often associated with DNA replication/repair events, may be important in the evolutionary sizing and shaping of genomes. A subset of these circles has been associated with human diseases (Autiero et al., 2002; Cohen et al., 1997).

The multi-copy 2-micron plasmid of *Saccahromyces cerevisiae*, present as 40-60 molecules per haploid genome content, is the most well characterized among yeast DNA plasmids (Broach & Volkert, 1991; Liu et al., 2014; Rizvi et al., 2018; Sau et al., 2019). Current evidence suggests that plasmid molecules are organized into clusters of 3-5 foci that act as units of segregation during cell division (Ghosh et al., 2007; Liu et al., 2013; Velmurugan et al., 2000). The plasmid genome has a bi-partite functional organization, namely, a replication-partitioning module and a copy number amplification module. The host replication machinery recognizes the plasmid origin, and duplicates each plasmid molecule once during S phase (Zakian et al., 1979). The plasmid partitioning system, comprised of the plasmid coded Rep1 and Rep2 proteins and the *cis*-acting *STB* locus, segregates the replicated molecules equally, or nearly so, to daughter cells (Liu et al., 2014; Rizvi et al., 2018; Sau et al., 2019). The amplification system, constituted by the plasmid coded Flp site-specific recombinase and its target *FRT* sites arranged in head-to-head orientation within the plasmid genome, compensates for copy number reduction resulting from rare missegregation events. Iterative copying of the plasmid template is triggered by a Flp-mediated recombination event coupled to bidirectional plasmid replication (Futcher, 1986; Petes & Williamson, 1994; Volkert & Broach, 1986). The expression of *FLP* is negatively regulated by a putative Rep1-Rep2 repressor, and positively by the plasmid coded Raf1 protein—an antagonist of the repressor (A. Rizvi et al., 2017; Murray et al., 1987; Reynolds et al., 1987; Som et al., 1988). The Flp protein levels and activity are controlled via post-translational modification by the host sumoylation machinery, which signals subsequent proteasome-mediated turnover of Flp (Chen et al., 2005; Ma et al., 2019; Xiong et al., 2009). Thus, mutually reinforcing regulatory mechanisms harbored by the plasmid and imposed by the host protect against any undue drop or runaway increase in plasmid population. Equal segregation and copy number optimization account for the evolutionary persistence of the 2-micron plasmid as a selfish DNA element that neither enhances nor significantly diminishes host fitness (Futcher & Cox, 1983; Mead et al., 1986).

The current model for 2-micron plasmid propagation, supported by several lines of circumstantial evidence, posits plasmid segregation in physical association with chromosomes (hitchhiking) (Liu et al., 2016; Liu et al., 2013; Mehta et al., 2002). Furthermore, plasmid sisters formed by replication of a single copy *STB*-reporter hitchhike on sister chromatids in a one-to-one fashion (Ghosh et al., 2007). This chromosome-like segregation is not only dependent on the Rep-*STB* system but also on plasmid replication. Apparently, the Rep-*STB* system, in conjunction with replication, directs the symmetric tethering of plasmid sisters to sister chromatids (Liu et al., 2016). In the absence of replication, the Rep-*STB*-assisted plasmid-chromosome tethering becomes random, and plasmid segregation follows the random assortment of chromosomes. Extrapolation of the single-copy plasmid behavior to the native multi-copy plasmid foci suggests a high-order organization within each focus for coordinating the replication of plasmid molecules and the symmetric attachment of the replicated copies to sister chromatids.

The orchestration of plasmid segregation by Rep1-Rep2-*STB* is mediated with the assistance of several host factors that interact with the partitioning system in a cell cycle dependent fashion (Cui et al., 2009; Ghosh et al., 2007; Ghosh et al., 2010; Hajra et al., 2006; Huang et al., 2011; Ma et al., 2012; Mehta et al., 2002; Mehta et al., 2005; Prajapati et al., 2017; Wong et al., 2002). Nearly all of these factors play a role in centromere (*CEN*)-mediated chromosome segregation as well. It has been speculated that the unusually short, and genetically defined, point-*CEN* of budding yeast chromosomes and the plasmid-*STB* have diverged from a common ancestor that once directed the segregation of both plasmids and chromosomes (Huang et al., 2011; Malik & Henikoff, 2009). Fluorescence tagged *STB*-reporter plasmids have suggested their tendency to be localized near centromeres and spindle pole bodies in mitotic cells (Velmurugan et al., 2000) whereas, in meiotic cells, their preferred localization is at or near telomeres (*TEL*s) (Sau et al., 2014). Plasmid-telomere association is dependent on the bouquet proteins Ndj1 and Csm1, which are meiosis-specific (Chu, 1998; Chua & Roeder, 1997; Conrad et al., 1997; Primig et al., 2000; Rabitsch et al., 2001). Subtle variations in the mode of plasmid-chromosome association and in the host factors responsible for this process may be necessitated by the two distinct cell cycle programs, even though the common hitchhiking theme is conserved in both cases. The role of the mitotic spindle, the spindle associated motor Kip1 and the microtubule plus end binding proteins, Bik1 and Bim1 in 2-micron plasmid segregation during mitosis (Cui et al., 2009; Mehta et al., 2005; Prajapati et al., 2017) suggests spindle-mediated recruitment of the 2-micron plasmid to chromosome sites in the vicinity of centromeres. The association of the *CEN*-specific histone H3 variant Cse4 with *STB* (Hajra et al., 2006), which has been additionally detected at centromere-like regions (CLRs) present near *CEN*s (Lefrancois et al., 2013), would also be consistent with the proximity between plasmid and *CEN*s.

The apparent differential plasmid localization on chromosomes in mitotic versus meiotic cells may, in principle, be reconciled by structural or organizational features common to centromeres and telomeres, and perhaps shared by other plasmid tethering sites on chromosomes. To address more definitively the still unresolved questions regarding chromosome locations utilized by the 2-micron plasmid for hitchhiking, we have now mapped the positions of a single-copy *STB*-reporter plasmid with respect to *CEN V* and *TEL V* (also *TEL V* and *TEL VII*) simultaneously. We find that the plasmid is located near *CEN* or *TEL* in ≥ 65% of the mitotic cells (≤ 0.5 μm), with a clear *TEL* preference over *CEN*. Interestingly, we observed that 2-micron plasmid segregation is impaired when the disjunction of strongly condensed loci such as telomeres or rDNA is blocked by Cdc14 inactivation. Consistent with this observation, induced missegregation of a highly condensin-dependent chromosome (Chr XII) has a stronger negative effect on equal plasmid segregation than the missegregation of a chromosome that is less dependent on condensin (Chr III). Genetic assays and chromatin immunoprecipitation (ChIP) analyses reveal the interaction of the condensin subunit Brn1 with the Rep-*STB* system, suggesting that condensin is an authentic plasmid partitioning factor appropriated from the host. Condensin requirement for plasmid segregation may be mandated by the need for multiple plasmid molecules within a focus to be condensed in order for them to co-segregate with a chromosome as one physical entity. Preferential localization at chromosome loci where condensin is enriched may reflect a plasmid strategy to satisfy this requirement. Alternatively, condensin acquisition by the clustered plasmid molecules, and presumably their condensation, may facilitate the tethering of plasmid foci to condensed chromosome locales. By hitchhiking on condensed, quiescent chromatin such as telomeres, the plasmid may minimize potential deleterious effects on chromosome function and on the host’s fitness.

## Results

### Strategy for mapping plasmid positions with respect to centromeres, telomeres and the spindle pole body at different stages of mitosis

Current evidence suggests that the 2-micron plasmid tends to localize preferentially near the centromere cluster and the spindle pole body in mitotic cells (Mehta et al., 2005), but migrates towards the nuclear periphery as diploid cells enter the meiotic program (Sau et al., 2014; Sau et al., 2015). These localizations were carried out primarily using fluorescence-tagged multi-copy *STB*-reporter plasmids (Mehta et al., 2005; Sau et al., 2014), and a nearly single-copy reporter plasmid in one instance (Cui et al., 2009). Visualization of the native 2-micron plasmid in mitotic cells using FISH also reveals a significantly higher number of plasmid foci unassociated with the nuclear membrane than those localizing with it (Heun et al., 2001a). Meiotic plasmid localization at the nuclear periphery increases from early meiosis to the pachytene stage, and requires the bouquet proteins Ndj1 and Csm4 (Sau et al., 2014). In pachytene chromosome spreads, over half the plasmid foci are situated on chromosomes, predominantly at or near telomeres (Sau et al., 2014). The sum of the available data, viewed in light of a shared chromosome-hitchhiking mechanism for plasmid segregation during mitosis as well as meiosis, is consistent with the following interpretation. The 2-micron plasmid exploits certain chromatin features characteristic of centromeres or telomeres, or regions proximal to them, for its chromosome localization. Plasmid presence distal to these loci may be accounted for by similar chromatin features present at a subset of other chromosome regions as well. Differential plasmid localization with respect to the nuclear periphery during mitosis and meiosis may reflect changes in chromatin organization, and/or altered spatial arrangements, of chromosome locales between mitotic and meiotic cells.

In order to gain further insights into the chromosome-associated segregation of the 2- micron plasmid, we sought to map plasmid positions in mitotic cells at higher resolution by simultaneously fluorescence-tagging chromosome V (Chr V) at the centromere and the telomere on the right arm. Furthermore, in order to avoid any imprecision posed by multiple plasmid foci, we employed a previously described *STB*-reporter plasmid (pSG1) present at one copy (and as a single focus) in ≥ 80% of the cells (Ghosh et al., 2007). The segregation behavior of this plasmid is quite similar to that of an *STB*-reporter plasmid excised from a chromosome by site-specific recombination at G1, and therefore has a copy number of exactly one prior to DNA replication (Liu et al., 2013). A unique advantage of pSG1, which also contains a *GAL* promoter-regulated centromere (*CEN*), is that it can be induced to behave as a *CEN*- plasmid, *STB*-plasmid or an *ARS*-plasmid (lacking partitioning activity) under appropriate experimental conditions (Figure 1- figure supplement 1). When cells harboring pSG1 are grown in medium with glucose as the carbon source (to repress the *GAL* promoter), it is referred to as pSG1-*CEN* to indicate an active centromere. The *CEN*-mediated partitioning function of pSG1- *CEN* is not affected by the presence or absence of the native 2-micron plasmid in the host cells ([Cir^+^] or [Cir^0^], respectively). When present in [Cir^+^] cells, and the carbon source is galactose, *CEN* is inactivated (by high level transcription from the *GAL* promoter), and the plasmid is referred to as pSG1*-STB* to denote its segregation by a functional Rep-*STB* partitioning system. When placed in [Cir^0^] cells, with galactose as the carbon source, the absence of the Rep proteins makes the plasmid behave as pSG1-*ARS* in segregation (Figure 1- figure supplement 1).

**Figure 1.**
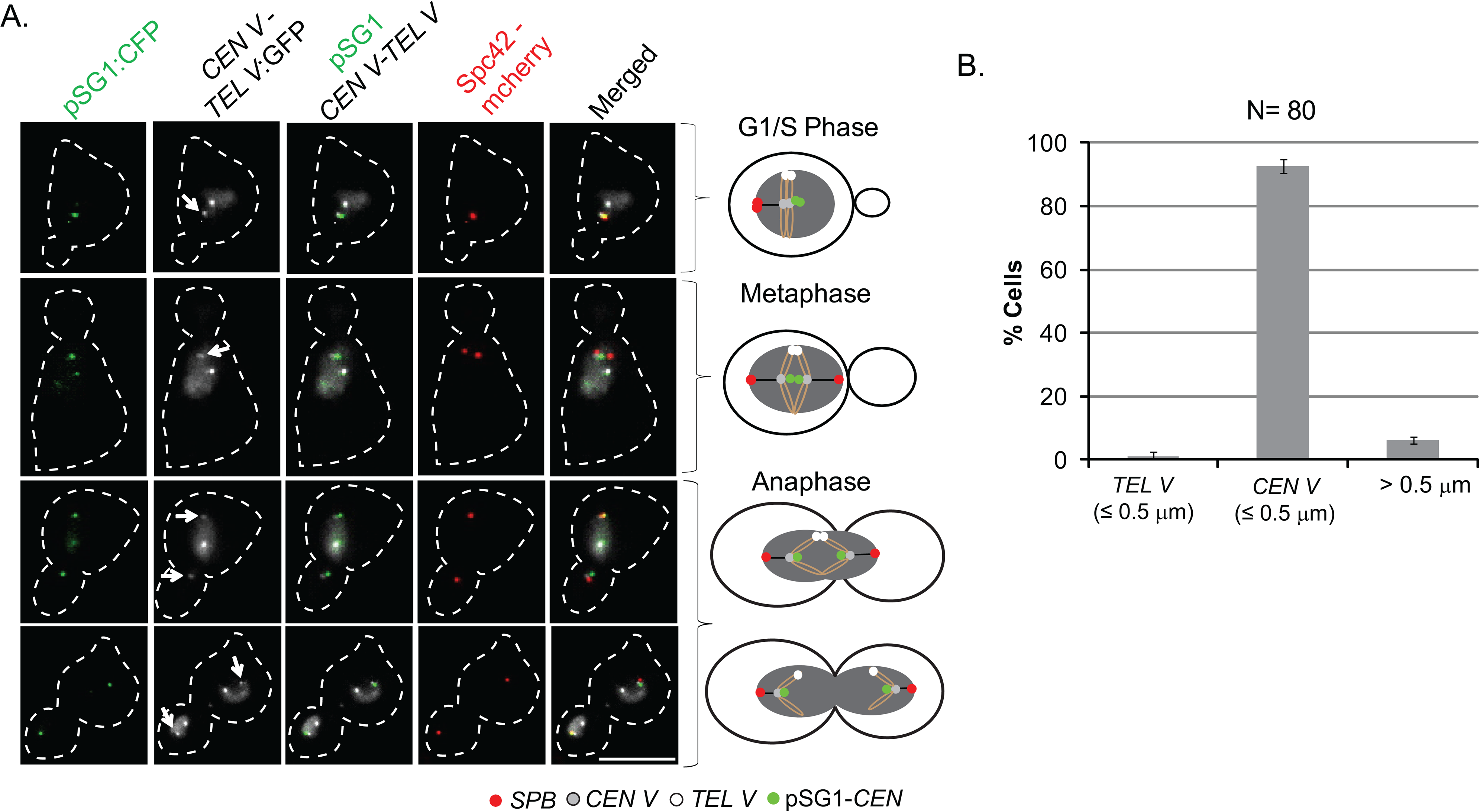
Localization of pSG1-*CEN* plasmid with respect to *CEN V* and *TEL V* during different stages of the cell cycle. **A**. The positions of pSG1-*CEN*, along with those of *CEN V* (indicated by arrows) and *TEL V*, were followed in [Cir^+^] cells grown in glucose (to keep the *CEN* harbored by the plasmid active). Representative images from fixed cells are shown in rows at the left, and summarized in the schematic diagrams at the right. **B**. pSG1 locations proximal to a reference locus (≤ 0.5 µm) or distal from either of the two reference loci (> 0.5 µm) are shown in the plots. ‘N’ represents total number of cells analyzed. Error bars denote the standard deviation of the mean acquired from three separate experiments. Bar, 5 µm.

We distinguished *CEN V* and *TEL V* in cells expressing TetR-GFP by their distinct fluorescence intensities conferred by [TetO]_224_ and [TetO]_448_ arrays inserted at *CEN V* and *TEL V* loci, respectively (Alexandru et al., 2001; Michaelis et al., 1997) (Figure 1- figure supplement 2). We verified that the two loci could be identified unambiguously at various stages of mitosis by marking their positions relative to the spindle pole body (SPB) tagged with Spc42-mCherry (Figure 1- figure supplement 2). In *S. cerevisiae*, centromeres remain congressed as a single cluster in close proximity to SPB during interphase as well as mitosis, while the rosette of three to six telomere clusters is positioned at the nuclear periphery distal to SPB in a Rabl-like pattern (Gotta et al., 1996; Guacci et al., 1997; Heun et al., 2001b; Jin et al., 2000; Jin et al., 1998; Kupiec, 2014; Taddei & Gasser, 2012). Consistent with this spatial organization, we observed partial overlap or close proximity of SPB to the fainter of the two GFP foci (*CEN V*), with the brighter GFP focus *(TEL V)* located away from it (Figure 1- figure supplement 2). These three nuclear landmarks provided a frame of reference for mapping the spatial positions of reporter plasmids.

### Mitotic localization of pSG1-*CEN* and pSG1-*STB* with respect to *CEN V* and *TEL V*

In order to follow plasmid localization in the nucleus, we introduced pSG1 into isogeneic [Cir^+^] and [Cir^0^] strains containing the marked *CEN V*, *TEL V* and SPB, and expressing cyan-LacI. Plasmid foci were visualized by repressor interaction with the plasmid-borne [LacO]_256_ array (Straight et al., 1996). As already noted, by using [Cir^+^] and [Cir^0^] backgrounds and manipulating the carbon source, we could induce pSG1 to behave as pSG1-*CEN*, pSG1-*STB* or pSG1-*ARS* (Figure 1/xref>- figure supplement 1, Ghosh et al., 2007).

The localization of pSG1-*CEN* ([Cir^+^]; glucose) at different stages of the mitotic cell cycle (Figure 1A, B) at or close to SPB (≤ 0.5 μm) was akin to that of chromosome centromeres (Figure 1- figure supplement 2), as expected. The spatial distribution of pSG1-*STB* ([Cir^+^]; galactose), however, was different (Figure 2 and 3). In G1/S (Figure 2A, B) and in metaphase (Figure 2C, D) cells, 40-45% of the cyan foci were proximal to the *TEL V* (≤ 0.5 μm), 20-25% were close to *CEN V* (≤ 0.5 μm) and the rest showed no spatial linkage to either *CEN V* or *TEL V* (> 0.5 μm) (Figure 2A-D).The pSG1-*STB* sisters were seen in metaphase cells as a single coalesced focus in the majority of cases (60-65%), as would be consistent with the bridging of plasmid sisters by the cohesin complex (Ghosh et al., 2010; Liu et al., 2013; Mehta et al., 2002) and/or their symmetric attachment to sister chromatids (Liu et al., 2016), which are also paired by cohesin. For the purpose of precise positioning, the cells (35-40%) showing plasmids with more than a single focus were excluded from the distance analysis. As the average copy number of pSG1 was only approximately one (Ghosh et al., 2007), not every cell in the population contained a single plasmid molecule prior to replication. At least a subset of the metaphase cells with two separated plasmid foci might indicate copy number deviation from unity, rather than a lack of pairing between plasmid sisters. Higher foci number seen in a very small fraction of cells could only be due to the presence of more than one plasmid per nucleus.

**Figure 2.**
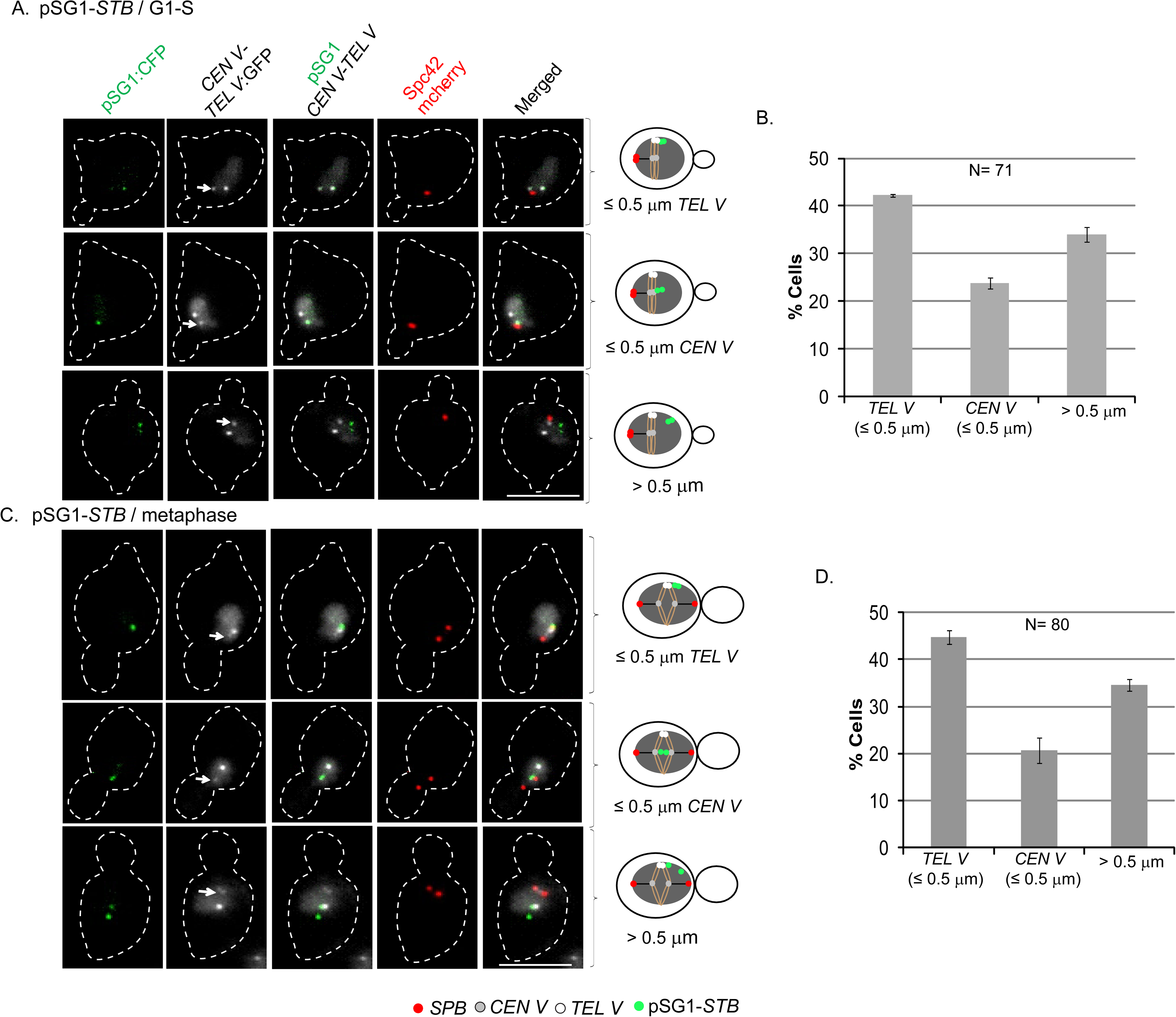
Preferred telomere-proximal localization of pSG1-*STB*. The localization of pSG1-*STB* ([Cir^+^]; galactose) with respect to *CEN V* and *TEL V* was scored in fixed cells at the G1/S (**A**) and metaphase (**C**) stages of the cell cycle. The types of plasmid localization with respect to reference loci represented by the rows of images at the left are codified in the corresponding schematic diagrams placed to the right of each row. The histogram plots in **B** and **D** are based on the data from **A** and **C**, respectively. ‘N’ represents total number of cells analyzed. Error bars denote the standard deviation of the mean acquired from three separate experiments. Bar, 5 µm.

**Figure 3.**
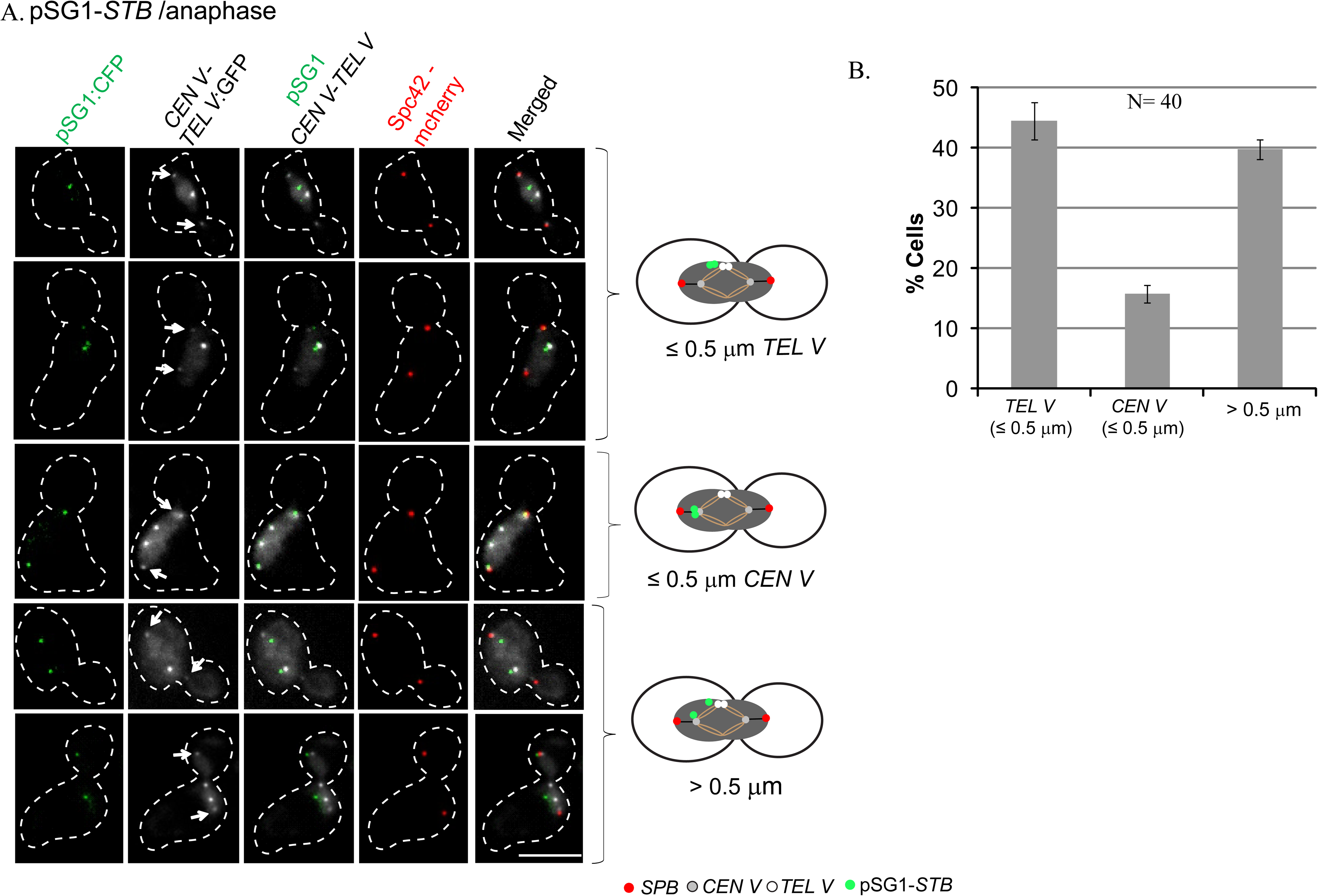
Preferential localization of pSG1-*STB* near telomeres revealed at the anaphase stage of mitosis. **A**. The representative images (rows at the left) depict a subset of early anaphase cells ([Cir^+^]; galactose grown) in which pSG1-*STB* was mapped with respect to well resolved *CEN V* (indicated by the arrows) and *TEL V*. The pSG1-*STB* positions were classified into three types as idealized by the schematic diagrams at the right. **B**. The plot shows the relative frequencies of the three types. ‘N’ represents total number of cells analyzed. Error bars denote the standard deviation of the mean acquired from two separate experiments. Bar, 5 µm.

As a more stringent test of the apparent tendency of pSG1-*STB* to remain telomere-proximal, we observed its localization in a selected subset of anaphase cells in which the *CEN V*s were well separated from each other, and from the unsegregated *TEL V*s that remained at the spindle mid-zone (Figure 3A). In this population, the fraction of cells showing pSG1-*STB* proximity to *TEL V* was ∼45%; *CEN V* proximity was seen in ∼15% (Figure 3B). For the pSG1- *ARS* plasmid, the proximal (≤ 0.5 μm from *CEN V* or *TEL V*) and distal (> 0.5 μm from *CEN V* and *TEL V*) locations of foci were roughly evenly divided among G1/S cells (Figure 2- figure supplement 1). As an *ARS*-plasmid does not physically associate with chromosomes, as revealed by chromosome spread assays (Liu et al., 2014; Mehta et al., 2002), the data in Figure 2- figure supplement 1 refer to the overall nuclear distribution of the plasmid and not its location on chromosomes.

Finally, we verified the mapping of plasmid positions by determining the three-dimensional disposition of pSG1 with respect to *CEN V* and *TEL V* in live cells. The dot plots of distances in ‘Figure 2- figure supplement 2’ were derived by defining the centroids of the plasmid and chromosome fluorescent foci from imaging a series of nuclear stacks (Figure 2- figure supplement 2A). The distribution of pSG1-*STB* showed a discernible bias towards telomere proximity (Figure 2- figure supplement 2B), while that of pSG1-*ARS* ([Cir^0^]; galactose) was essentially random (Figure 2- figure supplement 1 and 2C).

The expected centromere-proximal localization of pSG1-*CEN* was also evident (Figure 1, Figure 2- figure supplement 2D). The combined results from Figure 2 and Figure 3 suggest that telomeres and their proximal loci are more frequently habited by the 2-micron plasmid than centromeres and nearby regions. At all the stages of the cell cycle, a third population of cells showed no association of pSG1-*STB* with *CEN V* or *TEL V* locus (> 0.5 μm in Figure 2 and 3). It is likely that the 2-micron plasmid associates with other tethering sites distributed along chromosome arms, albeit at a lower frequency.

### Association of the 2-micron plasmid with *TEL VII*

Unlike centromeres, which are congressed into one tightly knit cluster in mitotic cells, telomeres form a rosette of 3-6 clusters (Kupiec, 2014; Taddei & Gasser, 2012). The premise of our assays (Figure 2 and Figure 3) is that *TEL V* is equally likely to associate with any of these clusters. In order to probe potential variations among telomeres in plasmid association, we assayed plasmid localization with respect to two telomeres, *TEL V* and *TEL VII* (on the right arm of Chr VII), differentially tagged by [TetO]_448_-[TetR-GFP] and [TetO]_∼50_-[TetR-GFP] fluorescence, respectively.

In the experimental strain *TEL V* was distinguished from *TEL VII* by the difference in their fluorescence intensities. The observed pSG1 association with *TEL V* was ∼48% and that with *TEL VII* was 24% (Figure 2- figure supplement 3). These results, while upholding the telomere preference of the 2-micron plasmid, suggest that all telomeres may not be equal in plasmid occupancy. Such differences might arise from differences in high-order chromatin organization and/or epigenetic features at or near individual telomeres. Some variability might also result if individual telomeres are not entirely bias-free in their associations during cluster formation. Furthermore, ‘proximity’ is somewhat broadly defined in our assays because of the resolution limits of fluorescence microscopy and the cut-off distance of 0.5 μm employed.

From the perspective of the hitchhiking model for segregation, the combined results from Figure 2 and Figure 3 suggest that telomere- and centromere-association together constitutes a major segregation route for the 2-micron plasmid. Furthermore, telomeres are better preferred as hitchhiking sites by the plasmid than centromeres. Thus, preferential plasmid segregation as a telomere appendage observed during meiosis (Sau et al., 2014) appears to hold for mitosis as well, a fact that was not appreciated from prior published results. While Rep1 and Rep2 proteins are essential for mediating plasmid-chromosome association in both mitosis and meiosis, the host proteins involved in plasmid linkage to telomeres are likely different between the two cell cycle programs (Sau et al., 2014).

### Blocking segregation of condensed chromosome loci causes *STB*-plasmid non-disjunction

Chromosome sites preferentially occupied by the condensin complex in budding yeast include telomere-proximal regions, which are organized as condensed, transcriptionally quiescent heterochromatin (Wang et al., 2005). Chromosome condensation is influenced by centromeres in their role as *cis*-acting mediators. They promote recruitment of condensin, cohesin and associated signaling molecules to pericentric regions, which have characteristic chromatin composition and organization (Biggins, 2013; Kruitwagen et al., 2018; Stephens et al., 2011; Yong-Gonzalez et al., 2007). Genome-wide mapping studies reveal quantitative differences in condensin occupancy along chromosome arms, as well as cell cycle dependent modulations in occupancy at individual loci (Wang et al., 2005). In mitotic yeast cells, in addition to sub-telomeric and pericentric regions, the rDNA gene cluster is also prominent in condensin enrichment. The non-random localization of an *STB*-plasmid in the nucleus, favoring the vicinity of telomeres and centromeres (Figure 2 and 3; Figure 2- figure supplement 2, Figure 2- figure supplement 3), prompted us to examine a potential role for condensin in 2-micron plasmid segregation, and to address the possible coupling between condensed chromosome loci and the plasmid in segregation.

Telomeres and rDNA differ subtly from chromosomes as a whole in their segregation mechanism. In addition to early anaphase cleavage of cohesin that holds together sister chromatids in pairs, these loci require a second ‘late’ condensin-dependent step to complete unlinking and segregation (Machin et al., 2016). The extra condensin recruitment is mediated by Cdc14 phosphatase, which is released from its sequestered state at the onset of anaphase by the FEAR (Fourteen Early Anaphase Release) network (Stegmeier & Amon, 2004; Sullivan et al., 2004). Upon inactivation of Cdc14, rDNA and telomeres are delayed in disjunction relative to the rest of the chromosomes (Clemente-Blanco et al., 2011). If the 2-micron plasmid preferentially utilizes condensed chromatin for its hitchhiking mode of segregation, lack of Cdc14 function is expected to increase the frequency of plasmid missegregation. We tested this prediction using a multi-copy *STB*-reporter plasmid pSV5-*STB* (Figure 4) that we had utilized in prior plasmid segregation assays (Mehta et al., 2002). The control reporter plasmids for this analysis were derivatives of pSV5 in which *STB* was either deleted (pSV6-*ARS*) or replaced by *CEN* (pSV7-*CEN*) (Figure 4- figure supplement 1 and 2).

**Figure 4.**
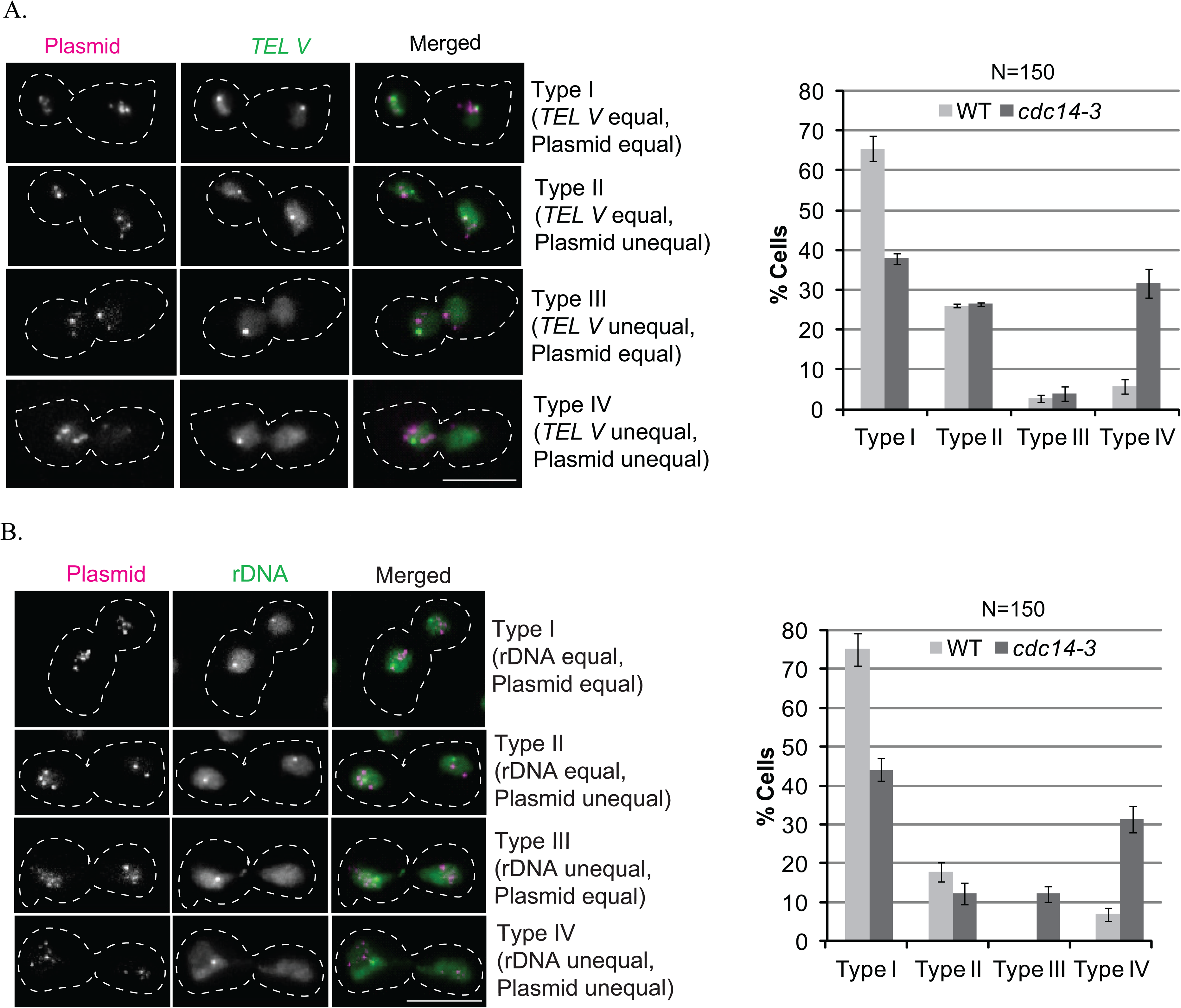
2-micron plasmid segregation when Cdc14 inactivation causes missegregation of telomeres or rDNA. **A**, **B**. The wild type and *cdc14-3* strains (otherwise isogenic) were arrested in G1 with α-factor at 23°C, and released from arrest at 33°C. Representative images of the four cell types analyzed are shown in the left panels (pSV5-*STB*, *TELV* in **A**; pSV5-*STB*, rDNA in **B**). The histogram plots for the quantitative estimates of the cell types are shown at the right. ‘N’ represents total number of cells analyzed. Error bars denote the standard deviation of the mean acquired from three separate experiments. Bar, 5 µm.

The pSV5 plasmid, fluorescence tagged by [LacO]-[CFP-LacI] interaction, is organized into 2-5 foci per nucleus. Prior experiments suggest that each plasmid focus is an independent unit in segregation (Velmurugan et al., 2000). As the number of plasmid molecules in individual foci is unknown, and occasionally foci tend to overlap, segregation estimates from counting foci numbers in mother and daughter nuclei are only semi-quantitative. Nevertheless, there is good agreement between data obtained from multi-copy and single-copy plasmid reporters in segregation assays (Ghosh et al., 2007; Liu et al., 2013; Velmurugan et al., 2000)

In the *cdc14-3* (Ts) strain at 33°C (non-permissive), a significant fraction of the cells showed non-disjunction of telomeres marked by [*TEL V*-LacO]-[GFP-LacI] (∼35%; Type III plus Type IV, Figure 4A) or the r-DNA locus marked by [rDNA–LacO]-[GFP-LacI] (∼40%; Type III plus Type IV, Figure 4B), as expected from prior studies (D’Amours et al., 2004). In the wild type strain, these loci segregated normally in ≥ 90% of the cells (Type I plus Type II, Figure 4A, B). The corresponding equal segregation of ∼70% for pSV5-*STB* (Type I plus Type III, Figure 4A, B;), which is consistent with prior results for multi-copy and single-copy reporter plasmids (Ghosh et al., 2007; Velmurugan et al., 2000), dropped to ∼40-50% in the *cdc14* mutant (Type I plus Type III, Figure 4A, B). Significantly, pSV5-*STB* and *TEL V* or rDNA were well correlated in their segregation, equal as well as unequal, in the wild type and mutant strains (Type I + Type IV >> Type II + Type III Figure 4A, B). In the majority of cells in which plasmid missegregation was coupled to the block in *TEL V* or rDNA segregation (Type IV, Figure 4A, B), the entire cluster of pSV5-*STB* foci, or a majority of the foci, was trapped in the mother or at the bud neck. In contrast to pSV5-*STB*, pSV6-*ARS* (Mehta et al., 2002), lacking *STB*, showed high missegregation in the wild type and mutant strains (Type II plus Type IV, Figure 4- figure supplement 1 A, B). On the other hand, pSV7-*CEN* segregated equally in most of the cells irrespective of the strain background (Type I and Type III, Figure 4- figure supplement 2 A, B). To further verify the apparent distinction between an *STB*-plasmid and an *ARS*- or a *CEN*- plasmid in segregation under Cdc14 inactivation, we followed the single copy reporter plasmids (pSG1-*STB*, pSG1-*CEN* and pSG1-*ARS*) with respect to Nop1 in the *cdc14* mutant strain (Figure 4- figure supplement 3). Consistent with the results from Figure 4, Figure 4- figure supplement 1 and Figure 4- figure supplement 2, pSG1-*STB* alone showed the correlation with Nop1 in segregation (Figure 4- figure supplement 3).

Thus, Cdc14 assisted segregation of *TEL V* or rDNA also impacts plasmid segregation directed by the Rep1-Rep2-*STB* system, but has no effect on *CEN* mediated segregation or segregation in the absence of a partitioning system. The correlation between *STB*-plasmid missegregation and the block in *TEL V* or rDNA segregation is consistent with condensed chromosome loci being preferred sites for hitchhiking by the 2-micron plasmid.

### Missegregation of chromosome XII causes missegregation of the 2-micron plasmid

Faithful chromosome segregation in yeast, as in all eukaryotes, requires chromosome condensation, which facilitates compaction of chromosome arms, unlinking of catenated sister chromatids, and their organization into functional segregation units (Hayes & Barilla., 2006). While condensation is easily visualized in the large chromosomes of plants and metazoans, more sensitive tools utilizing FRET or fluorescence quenching are needed to monitor condensation of the relatively short yeast chromosomes (Kruitwagen et al., 2018). However, condensation of the rDNA locus on chromosome XII (comprised of ∼150 copies of a 9.1 kbp repeat unit or 1.5 Mbp DNA) has been revealed by standard microscopy in conjunction with FISH (fluorescence in situ hybridization) (Lavoie et al., 2004). As already pointed out, segregation of the rDNA and telomere loci requires a Cdc14-dependent extra step of condensin recruitment and action. Interruption of this step perturbs the segregation of these loci and of the 2-micron plasmid, while chromosome segregation overall proceeds normally.

When gross missegregation of yeast chromosomes is induced by a conditional mutation, the majority of foci formed by a fluorescence-tagged multi-copy *STB*-plasmid segregates to the nuclear compartment containing the bulk of the chromosome mass (Mehta et al., 2002; Velmurugan et al., 2000). When the entire set of replicated sister chromatids is forced into either the mother or the daughter nucleus, a single-copy *STB*-plasmid is almost always localized in the chromosome-containing nucleus (Liu et al., 2016). These observations are consistent with plasmid hitchhiking sites being distributed on more than one (or perhaps all) chromosomes. However, the correlation between plasmid and Nop1 (rDNA) during segregation under Cdc14 inactivation suggests that missegregation of chromosome XII, housing the rDNA array and critically dependent on condensin for segregation, is likely to affect 2-micron plasmid segregation more severely than missegregation of a smaller chromosome with less stringent dependence on condensin. We tested this hypothesis by following plasmid behavior in cells missegregating either chromosome XII (the longest yeast chromosome) or chromosome III, which is roughly one-third as long.

The pSV5-*STB* plasmid was introduced into isogenic strains in which either *CEN III* or *CEN XII* could be inactivated conditionally by *GAL* promoter driven transcription through them (Reid et al., 2008). The expression of GFP-LacI and Nop1-RFP in these strains permitted plasmid segregation and chromosome XII segregation (by Nop1 as proxy) to be monitored simultaneously against chromosome segregation as a whole visualized by DAPI staining. Missegregation of chromosome XII, but not of chromosome III, resulted in a significant increase in plasmid missegregation (Figure 5A-C). Furthermore, in cells showing missegregation of both pSV5-*STB* and Nop1 (Type III; Figure 5A;), the majority of plasmid foci was present in the nucleus containing Nop1.

**Figure 5.**
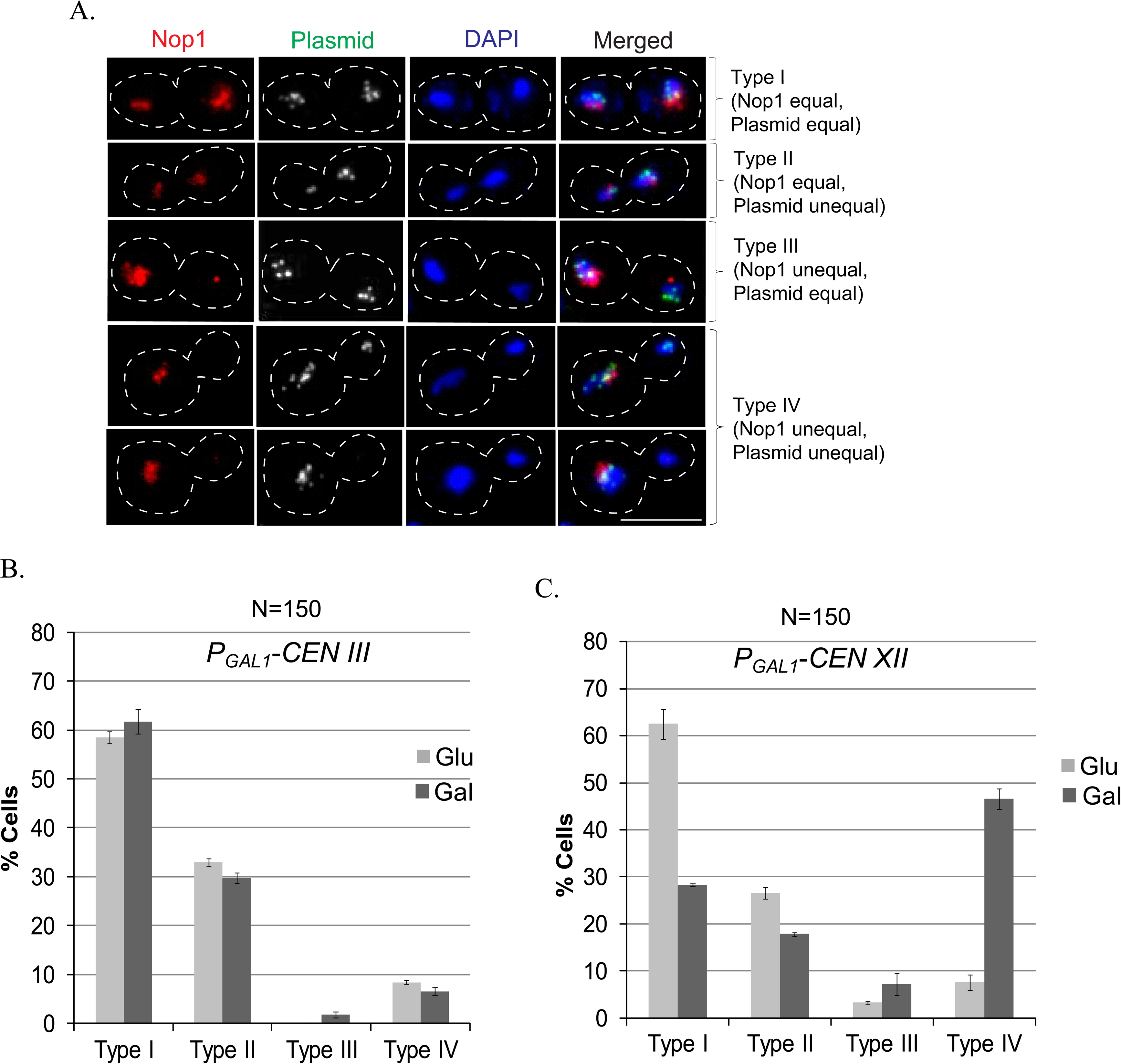
2-micron plasmid segregation in cells missegregating either chromosome III or chromosome XII. **A.** The experimental strains contained the *GAL1* promoter placed proximal to *CEN III* or *CEN XII* to drive high-level transcription through them under galactose induction. These strains were engineered to express GFP-LacI and Nop1-RFP for fluorescence tagging pSV5-*STB* and chromosome XII, respectively. Plasmid and chromosome segregations were assayed in anaphase cells under uninduced (glucose; *CEN III* and *CEN XII* active) or induced (galactose; *CEN III or CEN XII* inactive) conditions. The four segregation patterns analyzed (Types I-IV) are illustrated by representative cell images. **B, C.** The cell fractions showing Type I-IV segregation are plotted for *CEN III* inactivation (B) and *CEN XII* inactivation (C). ‘N’ represents total number of cells analyzed. Error bars denote the standard deviation of the mean acquired from three separate experiments. Bar, 5 µm.

The above results suggest that chromosome XII is a more preferred target for 2-micron plasmid association than chromosome III (and likely the other individual chromosomes). The size advantage of chromosome XII over chromosome III should have only minimal effect on plasmid association/segregation, as it would be more than offset by the fifteen normally segregating chromosomes in each of the assays.

### Absence of condensin function disrupts faithful *STB*-plasmid segregation

The bias towards condensed chromatin in the chromosome-association of the 2-micron plasmid suggests two possible roles, which need not be mutually exclusive, for the condensin complex in plasmid physiology. The multiple plasmid copies present within a chromosome associated plasmid focus, which appears to be the unit in segregation (Sau et al., 2019; Velmurugan et al., 2000), may need to be condensed as a prerequisite for segregation by hitchhiking. Or condensin may facilitate preferential plasmid association, and segregation in concert, with condensed chromosome loci. In support of the latter role, condensin is known to bridge DNA loci that are distal to each other on the genome. For example, it binds to tRNA genes dispersed throughout the chromosomes, and promotes their congregation into a cluster at the nucleolus (Haeusler et al., 2008). In order to probe potential condensin requirement for 2-micron plasmid segregation, we compared the rates of missegregation of *STB*-plasmids with those of *CEN*- or *ARS*-plasmids following inactivation of the condensin component Brn1.

At 36°C, the instability of the *STB*-plasmid (pRS422; *ADE2*) (Brachmann et al., 1998) was elevated ∼4-fold in the temperature sensitive *brn1-60* mutant (Ouspenski et al., 2000) over the wild type (Figure 6A). We also noted a comparable increase in the loss rate of the control *CEN*-plasmid (pRS412; *ADE2*) (Brachmann et al., 1998), but not an *ARS*-plasmid (YRp17, *TRP1*) (Botstein and Davis 1981) in the mutant background (Figure 6A). The *CEN*-plasmid instability is consistent with the previously demonstrated role of condensin in centromere function (Yong-Gonzalez et al., 2007). These stability results were further verified by assaying the segregation of fluorescence-tagged *STB*-reporter and *CEN*-reporter plasmids (pSV5-*STB* and pSV7-*CEN*, respectively) in the wild type and mutant strains (Figure 6B). A fluorescence-tagged *ARS*-plasmid (pSV6-*ARS*), by contrast, was not altered in its segregation by the loss of condensin function (Figure 6B).

**Figure 6.**
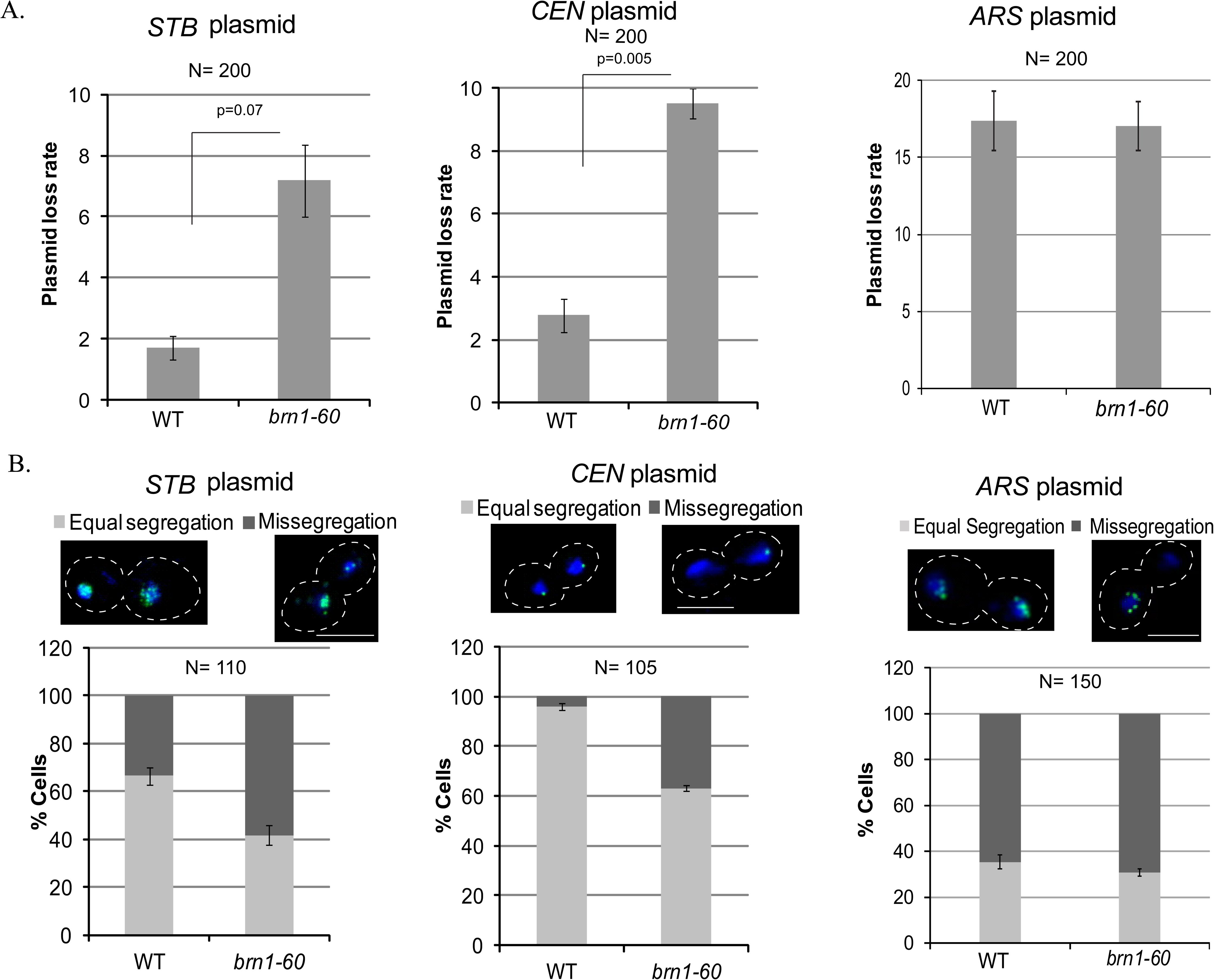
Missegregation of *STB*- and *CEN*-plasmids is increased in a condensin-defective strain relative to *ARS*-plasmids. **A.** Plasmid loss rates were estimated after 6-7 generations of non-selective growth at 36°C in the wild type and 4-5 generations in *brn1-60* strains (Materials and Methods). **B.** The segregation patterns of the fluorescence-tagged reporter plasmids were scored in the same strains as in (A). ‘N’ represents total number of colonies counted in (A) and cells analyzed in (B). Error bars denote the standard deviation from the mean values acquired from three independent experiments. p-values were calculated using *t*-test after comparing the values from WT and *brn1-60*. Bar, 5 µm.

The difference in the responses of *ARS*- versus *STB*- and *CEN*-plasmids to condensin inactivation suggests that the observed segregation results are unlikely to have resulted from the pleiotropic effects of condensin on chromatin organization and function. However, they do not address whether condensin plays similar or distinct roles in its contribution towards the segregation functions of *STB* and *CEN*.

### Brn1 interacts with the 2-micron plasmid partitioning system

Since condensin is required for chromosome segregation, and the 2-micron plasmid is dependent on chromosomes for its own segregation, it is difficult to decipher whether condensin’s effect on the plasmid is direct or indirect. Proximity of the plasmid to condensed chromosome loci may lead to incidental plasmid-condensin association, and likely plasmid condensation as a result. Alternatively, condensin acquisition by the plasmid (and perhaps condensation) may be functionally related to the localization of a plasmid focus at condensed chromatin and its chromosome associated segregation. One reasonable criterion for the direct involvement of condensin in 2-micron plasmid segregation would be its interaction with the Rep-*STB* system. Such interactions by host factors that assist plasmid partitioning have been demonstrated in previous studies through genetic interaction, chromatin immunoprecipitation (ChIP), and affinity enrichment-mass spectrometry assays (Cui et al., 2009; Hajra et al., 2006; Ma et al., 2013; Mehta et al., 2002; Prajapati et al., 2017). We have now subjected condensin to similar tests using the Brn1 subunit of the complex as its representative.

Interaction between *STB* and Brn1 was detected in a monohybrid assay, in which Brn1 fused to Gal4 transcriptional activation domain (Brn1-AD) increased the transcription of a *HIS3* reporter gene harboring *STB* as an upstream activator sequence (UAS) (Figure 7- figure supplement 1). This transcriptional activation was abolished in the absence of Rep1 and Rep2 ([Cir°] host) (Figure 7- figure supplement 1), suggesting that Brn1-*STB* interaction is mediated by Rep1 or Rep2, or by both. Consistent with Rep protein mediated condensin recruitment at *STB*, yeast dihybrid assays revealed the interaction of Brn1 with Rep1 and Rep2 (Figure 7- figure supplement 2).

**Figure 7.**
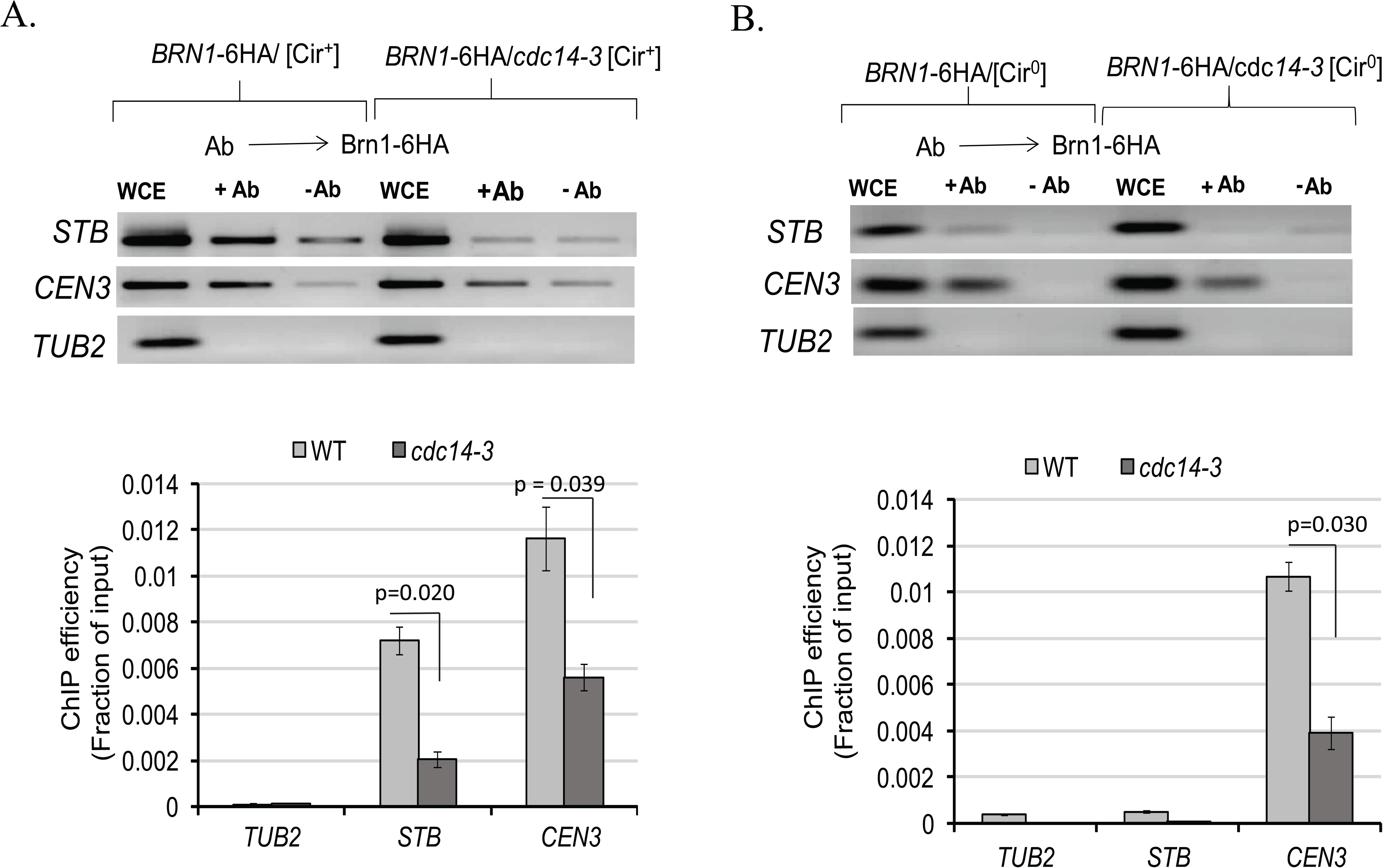
Brn1 associates with *STB* with the assistance of Rep proteins. **A, B.** ChIP analyses were performed using the indicated wild type and mutant strains (Cir^+^ or Cir^0^). Brn1-6HA was immunoprecipitated using an antibody to the HA-tag. The ΔCT values between an experimental sample and the input DNA were derived from qPCR data after correcting for primer efficiency/amplification factor (Verzijlbergen et al., 2014). ChIP efficiency was quantitated as the fraction of the input DNA that was immunoprecipitated [ε^(-ΔCT)], where ε is the amplification factor). The *TUB2* locus was used as a negative control (no Brn1 binding). Error bars denote the standard deviation from the mean values acquired from three independent experiments with minimum two technical replicates for each sample. p-values were calculated using *t*-test after comparing the values from WT and *cdc14-3*.

The results from the genetic assays were confirmed by the enrichment of *STB* DNA during chromatin immunoprecipitation (ChIP) in a [Cir^+^] strain using anti-HA antibodies directed to Brn1-6HA (Figure 7A). The authenticity of Brn1-*STB* association was verified by centromere (*CEN III*) enrichment, but not the tubulin locus *TUB2*, in the immunoprecipitate. *TUB2* has been designated as weak or negative in condensin localization (Wang et al., 2005). Furthermore, ChIP in a [Cir^0^] strain (lacking Rep1 and Rep2) failed to bring down *STB*, with little or no effect on *CEN III* (Figure 7B). Finally, consistent with the higher missegregation of *STB*- plasmids under Cdc14 inactivation (Figure 4, Figure 4- figure supplement 3), ChIP in the *cdc14-3* mutant revealed a nearly complete absence of condensin at *STB* (Figure 7A, B).

The Rep-protein-dependent association of condensin with *STB*, the loss of this association upon Cdc14 inactivation, and the rise in *STB*-plasmid instability upon Brn1 or Cdc14 inactivation argue for condensin being an authentic host coded 2-micron plasmid partitioning factor. The appropriation of the host’s DNA compaction and/or high-order DNA organization factors (condensin and cohesin, for example) by the 2-micron plasmid would be consistent with the evolutionary attunement of plasmid segregation to chromosome segregation.

## Discussion

We have characterized chromatin attributes exploited by the 2-micron plasmid for its hitchhiking mode of segregation during mitosis. Our findings suggest that condensed or condensin enriched chromatin and/or silent heterochromatin-like regions of chromosomes are preferred targets for plasmid-chromosome association. Furthermore, the condensin complex interacts with the plasmid partitioning system, and facilitates equal plasmid segregation.

### 2-micron plasmid localization on yeast chromosomes

The chromosome-hitchhiking model for 2-micron plasmid segregation (Liu et al., 2016; Liu et al., 2013; Rizvi et al., 2018; Sau et al., 2014) is based on a confluence of circumstantial, albeit persuasive, evidence. Strictly, plasmid association with a nuclear entity that segregates with the same characteristics as chromosomes cannot be ruled out. Direct demonstration of plasmid-chromosome association is impeded by the inability to resolve individual mitotic yeast chromosomes. Nevertheless, multi-copy and single-copy *STB*-reporter plasmids localize to yeast mitotic chromosome spreads in a Rep1-Rep2 dependent fashion, and colocalize with these proteins (Ghosh et al., 2006; Mehta et al., 2002). In mammalian cells, co-expressed Rep1 and Rep2 form merged foci on mitotic chromosomes collectively, and along the arms of individual chromosomes in metaphase spreads (Liu et al., 2016). Based on Rep1-Rep2 association with *STB* (Mehta et al., 2002; Scott-Drew & Murray, 1998; Velmurugan et al., 2000), and assuming the mammalian system to recapitulate the native yeast system, it follows that the 2-micron plasmid is tethered to yeast chromosomes with assistance from the Rep proteins. Meiotic yeast chromosome spreads, with considerably higher resolution than mitotic spreads, reveal *STB*-plasmid foci to be coalesced with a subset of Rep foci localized on chromosomes (Sau et al., 2014). The preferential plasmid localization at telomeres or subtelomeric regions in these spreads differs from the reported centromere proximity of plasmid in mitotic cells (Mehta et al., 2002). Higher accuracy mapping has now localized the 2-micron plasmid near *CEN* and *TEL* in a majority of the cell population, with a clear bias towards *TEL* (Figure 2 and Figure 3). In over a third of the population, the plasmid location is distal to *CEN* or *TEL*. It is possible that these ‘other’ chromosome sites that the plasmid tethers to share organizational features with *CEN* and *TEL*, for example, chromatin architecture and/or transcriptional quiescence.

### A role for chromatin organization in the selection of plasmid tethering sites?

*S. cerevisiae* lacks the canonical eukaryotic machinery for silenced heterochromatin assembly (Briggs et al., 2001; Grunstein & Gasser, 2013), but is still capable of transcriptional repression through specialized chromatin architecture. Silencing at telomeres, rDNA and the *HML*/*HMR* loci requires one or more components of the Sir protein complex (Sir1-4) along with other companion proteins that are shared among, or unique to, individual loci (Huang et al., 2002; Kueng et al., 2013). The atypical, genetically defined point centromeres of *S. cerevisiae* chromosomes do not conform to the classical definition of heterochromatin observed at epigenetically determined regional centromeres. Nevertheless, Sir1 is associated with *CEN*, and contributes to mitotic chromosome stability (Sharp et al., 2003), which can be modulated by driving transcription through *CEN* (Hill & Bloom, 1989; Ohkuni & Kitagawa, 2012). By a more inclusive definition, ‘heterochromatin’ of *S. cerevisiae* encompasses not only transcriptionally inactive chromatin but also functional *CEN* chromatin. The plasmid partitioning system may recognize tethering sites, including the preferred *TEL* and *CEN*, by a chromatin signature shared within this broad class of chromosome regions.

The *S. cerevisiae* point centromere, despite being genetically specified, manifests a certain degree of epigenetic character. Established *CEN*s are functionally propagated through many generations in strains carrying kinetochore mutations that block *de novo CEN* establishment (Mythreye & Bloom, 2003). By one postulate, centromeres originated from telomeres in parallel with the microtubule based cytoskeletal segregation machinery in response to the formation of multiple linear chromosomes from an ancestral single circular genophore (Villasante et al., 2007). *CEN* and *TEL* do share organizational or functional features in a limited set of biological contexts. In the fission yeast, centromere-centrosome contact can replace telomere-centrosome contact to support spindle formation and successful meiosis (Fennell et al., 2015). In fruit fly oocytes, the clustering of centromeres resembles bouquet formation by telomeres, which is emblematic of eukaryotic meiosis in general (Takeo et al., 2011). Both *TEL*s and *CEN*s being tethering sites for the native yeast plasmid would be consistent with some epigenetic mark that was conserved during their separate evolutionary trajectories, which might also be retained within *CEN*-adjacent centromere like regions (*CLR*s) (Lefrancois et al., 2013).

### 2-micron plasmid missegregation upon Cdc14 inactivation

The functional relevance of telomere proximity of the 2-micron plasmid on chromosomes is upheld by the increase in plasmid missegregation when telomere (and rDNA) disjunction is impeded by inactivating Cdc14. However, total missegregation of all plasmid foci in tandem with *TEL* or rDNA (represented by Nop1) is rarely seen (Figure 4). This is the expected outcome if plasmid tethering sites were present at chromosome locales other than *TEL*s, including *CEN*- proximal regions as revealed in the localization assays. According to a prior study, when the entire set of chromosomes is confined to either the mother or daughter nucleus, there is near perfect correlation between an *STB*-plasmid and chromosomes in their localization (Liu et al., 2016). Taken together, these results support a non-random distribution of plasmid tethering sites on chromosomes, *TEL*s or nearby loci being high density or high affinity sites. Cdc14 inactivation has pleiotropic consequences, one of which is compromised spindle integrity (Khmelinskii & Schiebel, 2008; Marston et al., 2003). The mitotic spindle and spindle associated proteins play a role, even if indirect, in 2-micron plasmid segregation (Mehta et al., 2002; Mehta et al., 2005; Prajapati et al., 2017). Hence, potential contribution of spindle defect to plasmid missegregation under loss of Cdc14 function cannot be ruled out. However, such an effect is likely modest, as overall chromosome segregation (revealed by DAPI staining) appears to be normal in our assays. Furthermore, *STB* plasmid missegregation can also be induced by inactivating the condensin component Brn1, which is not known to be involved in spindle function.

### A role for the condensin complex in 2-micron plasmid segregation

As noted, Cdc14-promoted condensin recruitment late in the cell cycle at rDNA and *TEL*s is a pre-requisite for their proper segregation (D’Amours et al., 2004; Sullivan et al., 2004). The possibility that this instalment of condensin resolves rDNA sisters independent of chromosome condensation and Topo II activity has been raised (D’Amours et al., 2004). However, full condensation of rDNA, approximately equal to that of condensed euchromatin, is completed only in anaphase (Sullivan et al., 2004). Resolution of the interlinks between rDNA sisters by recruitment of Topo II in a condensin dependent step, or remodeling of the interlinks by condensin to a configuration favored by Topo II, has not been ruled out (Charbin et al., 2014; Sullivan et al., 2004). The increased missegregation of the 2-micron plasmid in a condensin mutant (Figure 6) may result from a direct effect of condensin on the plasmid itself, or an indirect effect manifested through chromosomes on which it hitches a ride.

By being a chromosome appendage, the plasmid becomes subjected to the same physical and conformational constraints that a chromosome experiences during segregation. Chromosome condensation and plasmid condensation may thus be coincidental events. However, the interaction of Brn1 with the Rep proteins and its localization at *STB* with Rep protein assistance (Figure 7, Figure 7- figure supplement 1 and 2) would be consistent with a more direct role for condensin in plasmid segregation. Interestingly, Brn1 association with *STB* is reduced by Cdc14 inactivation (Figure 7). Perhaps Cdc14 dependent condensin assembly is shared by repeated loci that include rDNA and telomeres on chromosomes and the extra-chromosomal (but chromosome associated) 2-micron plasmid. Cohesin-mediated pairing of plasmid sisters (Ghosh et al., 2010; Mehta et al., 2002), in conjunction with possible condensin-assisted DNA compaction, may promote the organization of replicated plasmid molecules into two equally populated sister clusters that tether symmetrically to sister chromatids. Such a mechanism is consistent with the segregation pattern observed for single-copy *STB* plasmids whose sister copies segregate one-to-one in most cells (Ghosh et al., 2007; Liu et al., 2016; Liu et al., 2014). Condensin associated with *STB* may also serve to recruit Topo II to the plasmid and/or assist decatenation of plasmid sisters. Topo II inactivation causes an accumulation of replicated 2-micron plasmids as catenanes (DiNardo et al., 1984).

### Hitchhiking on host chromosomes by selfish DNA elements: choice of tethering sites

The 2-micron plasmid resembles viral episomes in infected mammalian cells in hitchhiking on chromosomes (Kanda et al., 2007; Nanbo et al., 2007; Coursey & McBride, 2019; Sau et al., 2019) even though their respective hosts are separated in evolution by ∼1.5 billion years. Given the high fidelity of chromosome dissemination to daughter cells, it is not surprising that the yeast plasmid and mammalian viruses have converged on the common strategy of exploiting the host’s chromosome segregation machinery for self-preservation. The telomere- and centromere-proximal localization of the 2-micron plasmid in mitotic cells suggest a propensity for plasmid tethering sites to be embedded in chromosome regions sparsely populated by genes or those organized into silent chromatin. Such localization would not only minimize fitness costs to the host but would also be advantageous to the plasmid. On the one hand, the host would be largely protected against perturbations to gene functions; on the other, the probability of plasmid dislodgement from the chromosome due to gene activity would be small.

Strikingly, the tethering sites on mammalian chromosomes for viral episomes also appear to be enriched for condensed chromatin (Aydin & Schelhaas, 2016). The heterochromatin regions that the episomes localize to include pericentromeric, peritelomeric and rDNA loci (Griffiths & Whitehouse, 2007; Kelley-Clarke et al., 2007; Krithivas et al., 2002; Oliveira et al., 2006; Poddar et al., 2009; Sekhar et al., 2010). A recent genome-wide Hi-C analysis has identified repressive chromatin as preferred tethering sites for Epstein-Barr virus (Moquin et al., 2017). Thus, the logic of associating with chromosomes for long-term persistence with as little disturbance to host genome structure and homeostasis as possible appears to be preserved among highly diverse eukaryotic selfish DNA elements.

## Materials and methods

### Yeast strains and plasmids

The yeast strains used in this study (W303 background) and their relevant genotypes are listed in Table S1. Strains containing or lacking the native 2-micron plasmid are referred to as [Cir^+^] or [Cir^0^], respectively. The [Cir^+^] designation does not include [Cir^0^] strains transformed with 2- micron derived or other artificial plasmid constructs. In the strains for chromatin immunoprecipitation (ChIP) assays, the *BRN1* locus on chromosome II was tagged with 6-HA by inserting a PCR amplified DNA fragment containing the epitope tag via homologous recombination (Janke et al., 2004). Gene fusion and expression of the tagged proteins were verified by PCR using total extracted DNA as template, and by immunostaining chromatin spreads with HA-antibody, respectively

The plasmid constructs employed in this work and their salient features are summarized in Table S1. The plasmid pRS402CFP-LacI (*ADE2*) was obtained by replacing the coding region for GFP in pRS402GFP-LacI (Velmurugan et al., 2000) with that of CFP.

### Tagging *TEL VII* with TetO

CRISPR (Cas9-sgRNA) technology was used to tag *TEL VII* with a [TetO]_n_ array in a strain expressing [TetR-GFP] and containing [TetO]_448_ inserted at *TEL V* (Ma et al., 2019). The PCR amplified DNA containing [TetO]_n_ (obtained from pRS306-[TetO]_224_ as template) flanked by chromosome homology at either end was inserted between *IMA1* and *MAL13* genes on the right arm of Chr VII (∼22 kbp from the end). Viable colonies obtained following transformation were screened by fluorescence microscopy to identify successful integrants. Based on the sizes of PCR products amplified using isolated total DNA, the number of [TetO] copies was < 50, and varied in individual transformants. One transformant showing two green fluorescent foci (bright *TEL V* and faint *TEL VII*) in most individual cells was saved for further analysis.

### Cell cycle synchronization

Cells were arrested in G1/S with α-factor, and released into the cell cycle by washing off the pheromone. Reporter plasmids were localized in the arrested cells and following their release. Cells harvested at intervals during cell cycle progression were examined by microscopy. Metaphase and anaphase cells were identified by their characteristic morphological features, as described previously (Cui et al., 2009; Ghosh et al., 2007).

### Plasmid stability assays

Cultures of transformant colonies containing a reporter plasmid were grown overnight in selective liquid medium, and were diluted into YPDA medium (non-selective). The inocula were grown for ‘n’ generations in non-selective media (n varied from 5 to 10, depending on the experiment). The fractions of plasmid containing cells in the population at the start (f_0_) and at the end (f_n_) of growth in the non-selective media were estimated by plating equal aliquots of the cultures on selective and non-selective plates. The plasmid loss rate per generation (%) was estimated as ‘i’ = (1/n) x ln(f_0_/f_n_) (Murray & Cesareni, 1986). The average values were plotted in a graph where error bars represented the standard deviation from the mean values acquired from three independent experiments. p-values were calculated using *t*-test (assuming unequal variances) after comparing the values from WT and *cdc14-3*.

### Plasmid segregation under induced missegregation of chromosomes

For inducing chromosome III or XII to missegregate in galactose medium, the *GAL1* promoter was inserted immediately upstream of *CEN III* or *CEN XII* as described previously (Reid et al., 2008). Cells harboring the *GAL-CEN* cassette were grown till mid-log phase in SC (-Leu, -Trp, raffinose) medium at 30°C before being transferred to SC (-Leu,-Trp) medium supplemented with either 2% galactose or 2% glucose for 5 hr at 30°C. Cells were harvested, stained with DAPI, and reporter plasmids ([LacO]-[GFP-LacI]) and rDNA (DsRed-Nop1) were observed by fluorescence microscopy to follow the segregation pattern of plasmid with respect to rDNA. The average values were plotted in a graph where error bars represented the standard deviation from the mean values acquired from three independent experiments.

### Monohybrid and di-hybrid assays

The monohybrid assay was performed as described earlier (Prajapati et al., 2017) to detect the interaction between *STB* locus and Brn1. The experimental strain was engineered to express *BRN1-*AD (AD = *GAL4* activation domain), and harbored *STB* placed upstream of the *HIS3* reporter locus as its UAS. Enhanced *HIS3* transcription as a result of Brn1-*STB* interaction (revealed by growth on medium lacking histidine) was assayed by inhibiting the His3 activity resulting from basal expression with 40 mM 3-AT (3-amino-1,2,4-triazole).

In the strain for di-hybrid assays (Finley et al., 1996), the bait protein was expressed as a fusion to LexA DNA binding domain, and LexO was integrated upstream of the *LEU2* reporter gene. A prey protein to be tested was fused to the B42 transcription activation domain, and was expressed by galactose induction from the *GAL1* promoter. The interaction between the bait and its putative prey was certified by colony growth in the absence of leucine. A positive response was verified by reciprocally swapping the bait and prey in the fusion proteins. Expression of the activation domain alone (without fusion to the bait) provided the negative control. Physical interaction between Rep1 and Rep2 was used as positive control for the assay.

### Chromatin immunoprecipitation assays

Chromatin immunoprecipitation (ChIP) assays were performed as described previously (Prajapati et al., 2018; Verzijlbergen et al., 2014). Brn1 fused to 6XHA at the carboxyl-terminus was immunoprecipitated using a polyclonal antibody to HA (Rabbit polyclonal HA11, Abcam, UK). The specificity of the association was verified using a chromosomal locus ‘*TUB2*’ designated to be weak or negative in condensin association (Wang et al., 2005). The relative amounts of immunoprecipitated DNA were quantitated using qPCR (Prajapati et al., 2018). Corrections were applied to account for primer efficiencies below 100% (amplification factors < 2.0) using standardization graphs of CT values against dilutions of the input DNA. The amplification factor ε was estimated as 10^(−1/slope) of the regression line, and the primer efficiency E (%) as {[10^(−1/slope)] – 1} x 100. The fraction of immunoprecipitated DNA in a ChIP sample relative to the input DNA was calculated as ε^∧^(−ΔCT), ΔCT= CT (ChIP) – [CT (Input) – log_ε_(Input dilution factor)]. The average values were plotted in a graph where error bars represented the standard deviation from the mean values acquired from three independent experiments. p-values were calculated using *t*-test (assuming unequal variances) after comparing the values from WT and *cdc14-3*.

### Fluorescence microscopy and distance measurement

Fluorescence microscopy was performed in live or mildly fixed cells. Fixing was performed in *p*- formaldehyde (4% v/v) by incubating on ice for 5-10 min. After two washes with 0.1 M phosphate buffer (pH 7.4), cells were imaged using a Zeiss Axio Observer Z1 fluorescence microscope and the Axiovision software. Images were acquired using the z-stack mode, with a 0.20 µm interval between two successive planes (Bystricky et al., 2005). They were opened in Imaris software (Bitplane, Imaris 8.0.2), and ‘slice tool’ was utilized to measure the spacing between the centroids of two fluorescent foci (Mittal et al., 2020). The distances were grouped into two types, ≤ 0.5 μm and > 0.5 μm. The average values were plotted in a graph where error bars represented the standard deviation from the mean values acquired from three independent experiments.

## Acknowledgements

This work was funded by DST (SB/SO/BB-0125/2013) and DBT (BT/PR13909/BRB/10/1432/2015) Govt. of India grants to SKG and NSF grants (MCB-1049925 and MCB-1949821) and a Robert F Welch Foundation award (F-1274) to MJ. HKP, DK and PM were supported by MHRD (10I30006), DBT (DBT/2018/IIT-B/1059) and UGC (17-06/2012(i) EU-V) fellowships, respectively. We acknowledge the central instrumentation facility of IIT Bombay for providing the microscope.

## Abbreviations

*CEN*: Centromere
*TEL*: Telomere
3-AT: 3-Amino-1,2,4-triazole

---

**Figure 1- figure supplement 1.**
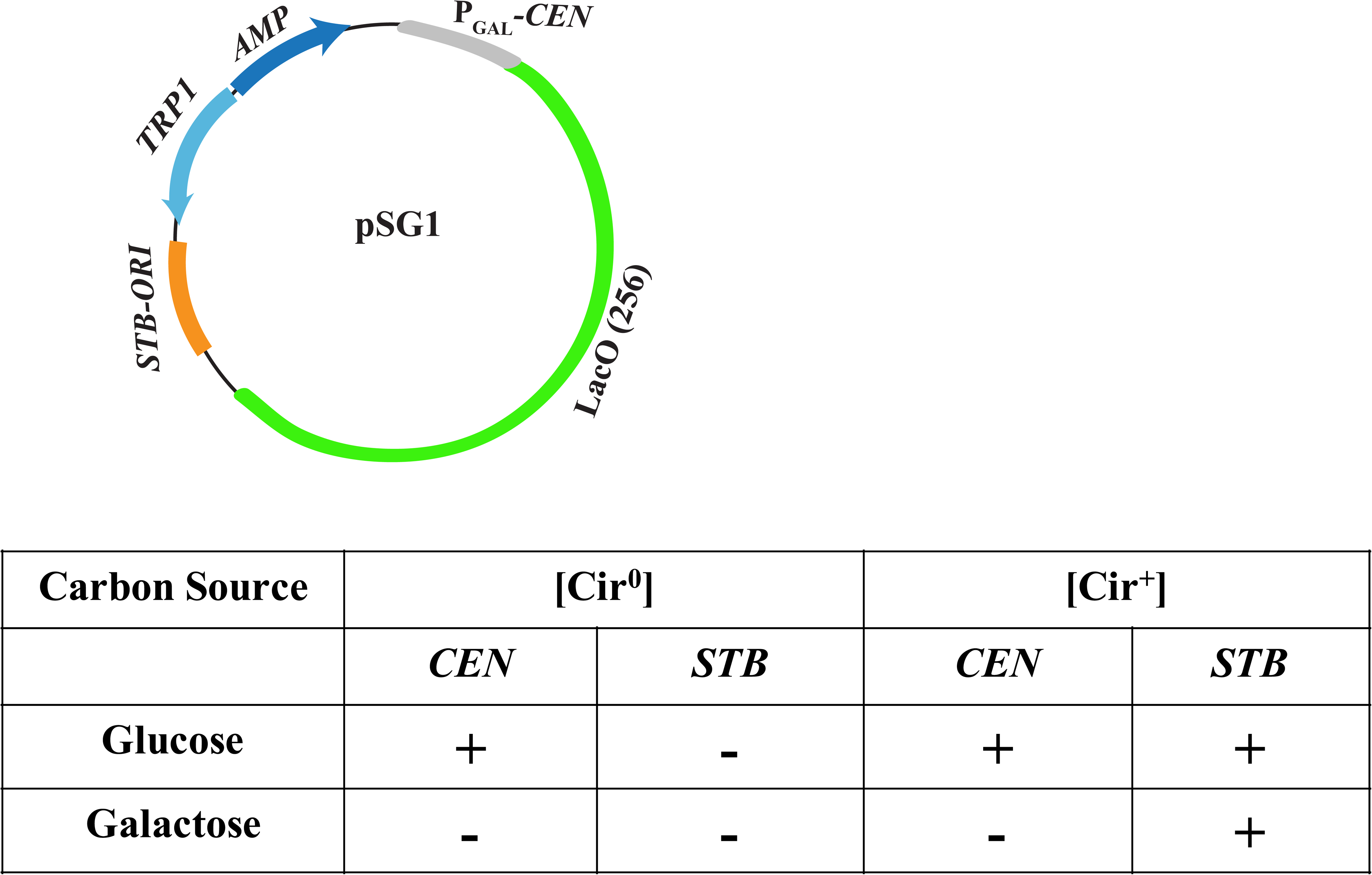
The functional states of *CEN* and *STB* in plasmid pSG1 under distinct experimental contexts. In the pSG1 plasmid, schematically diagrammed at the top, the centromere (*CEN*) is placed immediately downstream of the *GAL* promoter. The segregation status of the plasmid in a [Cir^0^] or [Cir^+^] host strain under glucose or galactose as the carbon source is tabulated below. *CEN* is active in both hosts when the promoter is turned off by glucose repression. It is inactivated by galactose-induced transcription. The Rep1 and Rep2 proteins provided by the native 2-micron circle of the [Cir^+^] strain keep *STB* active, regardless of the carbon source. In the [Cir^0^] strain, lacking Rep1 and Rep2, *STB* is inactive. In the [Cir^0^]/galactose context, neither *CEN* nor *STB* is active. As a result, pSG1 segregates as an *ARS* plasmid. The active and inactive states of *CEN* or *STB* are indicated by ‘+’ and ‘-‘, respectively.

**Figure 1- figure supplement 2.**
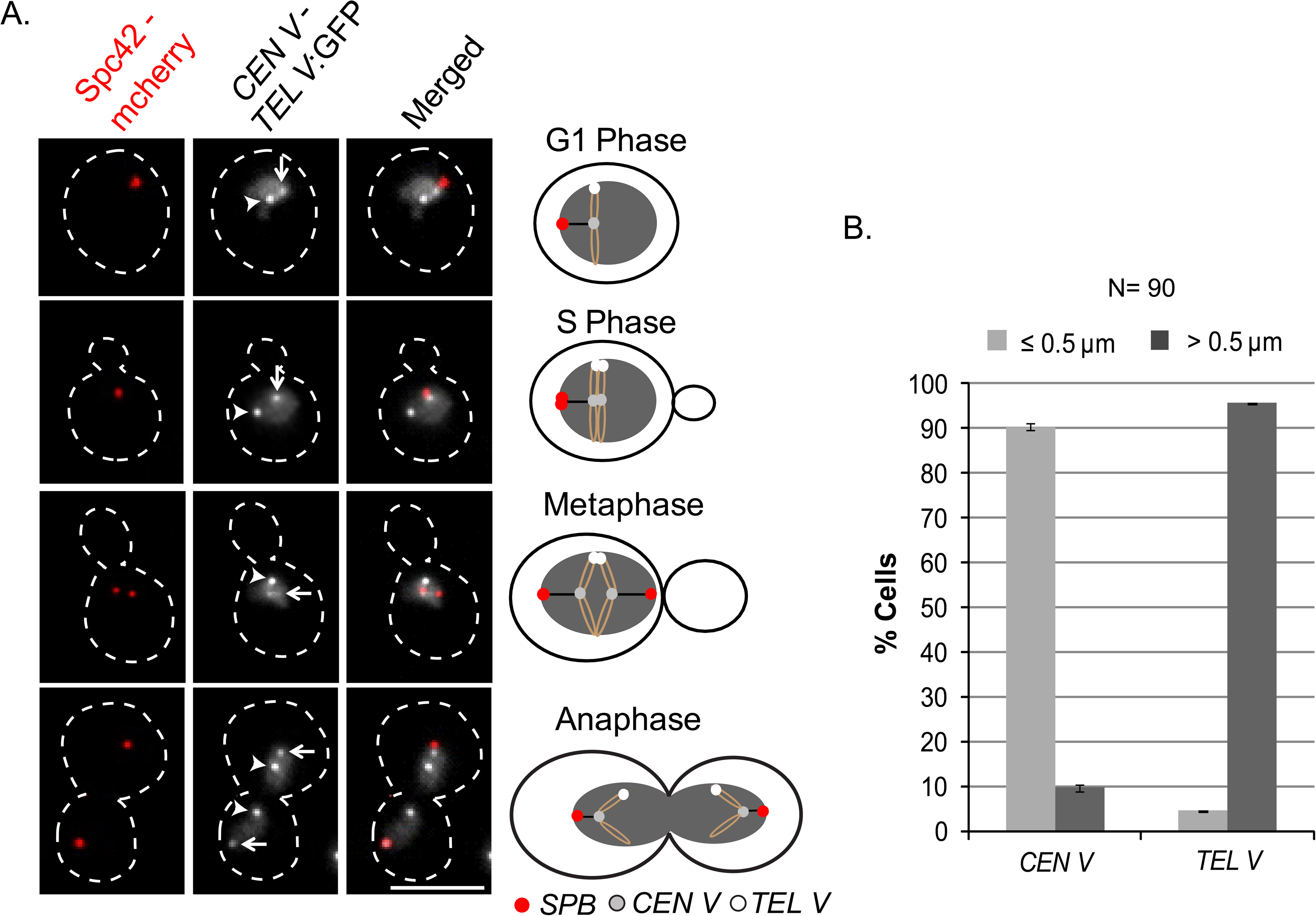
The localization patterns of SPB, *CEN V* and *TEL V* during different stages of the mitotic cell cycle. In the experimental strain, SPB was marked with red fluorescence using Spc42 fused to m- herry. *CEN V* and *TEL V* were tagged by green fluorescence via [TetO]_n_-[TetR-GFP] interaction. These two loci were distinguished by their differential brightness ([TetO]_224_ at *CEN* and [TetO]_448_ at *TEL*). **A.** The representative images from fixed cells (left) and their schematic illustrations (right) denote the positions of the three nuclear landmarks at different stages of the cell cycle. The arrows and arrowheads point to *CEN V* and *TELV*, respectively. Bar, 5 µm. **B.** The histograms show distance measurements of *CEN V* and *TEL V* from SPBs that are divided into two categories (≤ 0.5 µm and > 0.5 µm). ‘N’ represents total number of cells analyzed. Error bars denote the standard deviation from the mean values acquired from three independent experiments.

**Figure 2- figure supplement 1.**
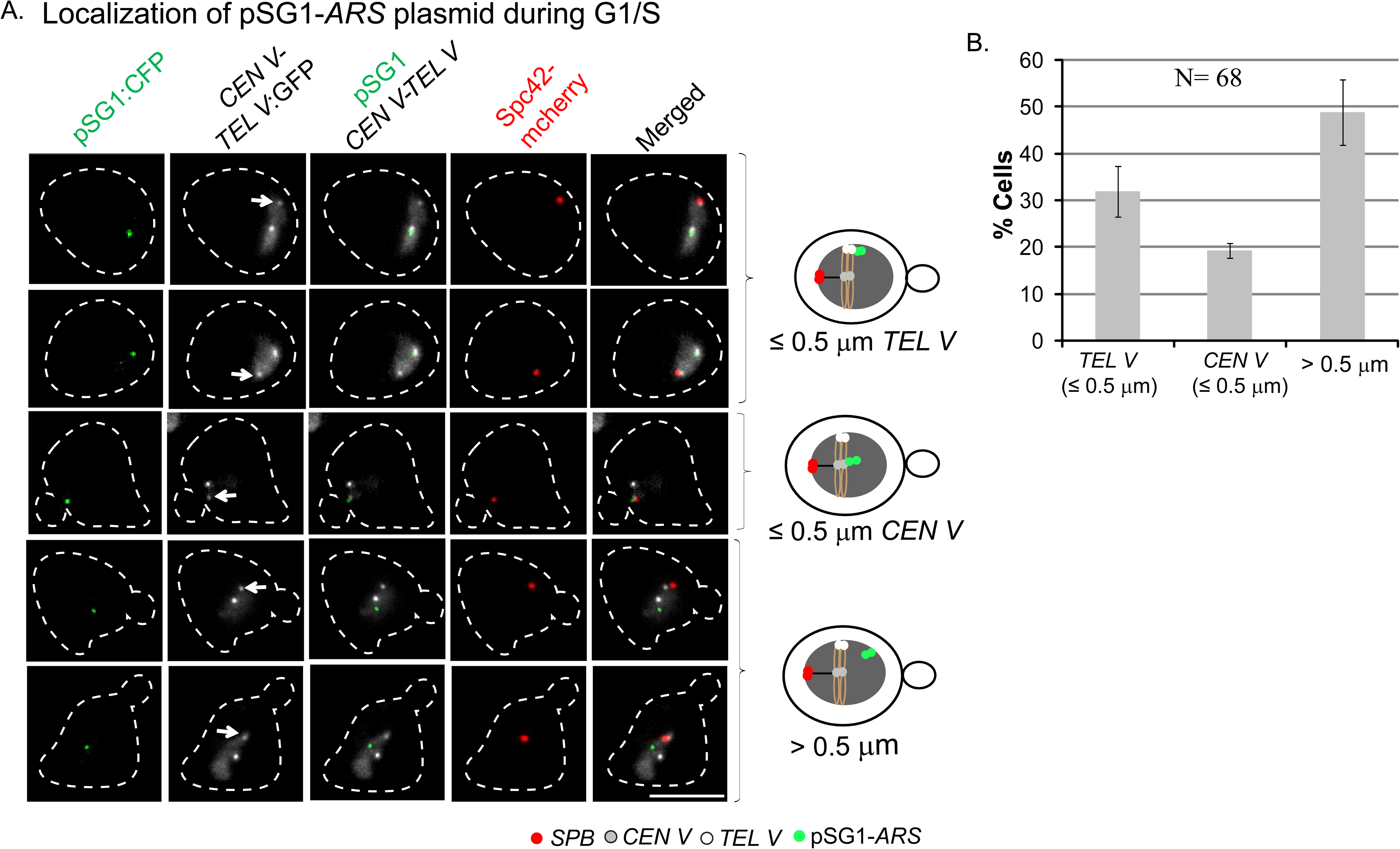
Localization of pSG1-ARS in the nuclei of mitotic cells at the G1/S stage. **A**. The cell images (left) and the corresponding schematic diagrams (right) represent the three localization patterns of pSG1-*ARS* assayed in [Cir^0^] cells grown in galactose. **B**. The relative abundance of each pattern is shown in the histogram plot. ‘N’ represents total number of cells analyzed. Error bars denote the standard deviation from the mean values acquired from three independent experiments.

**Figure 2- figure supplement 2.**
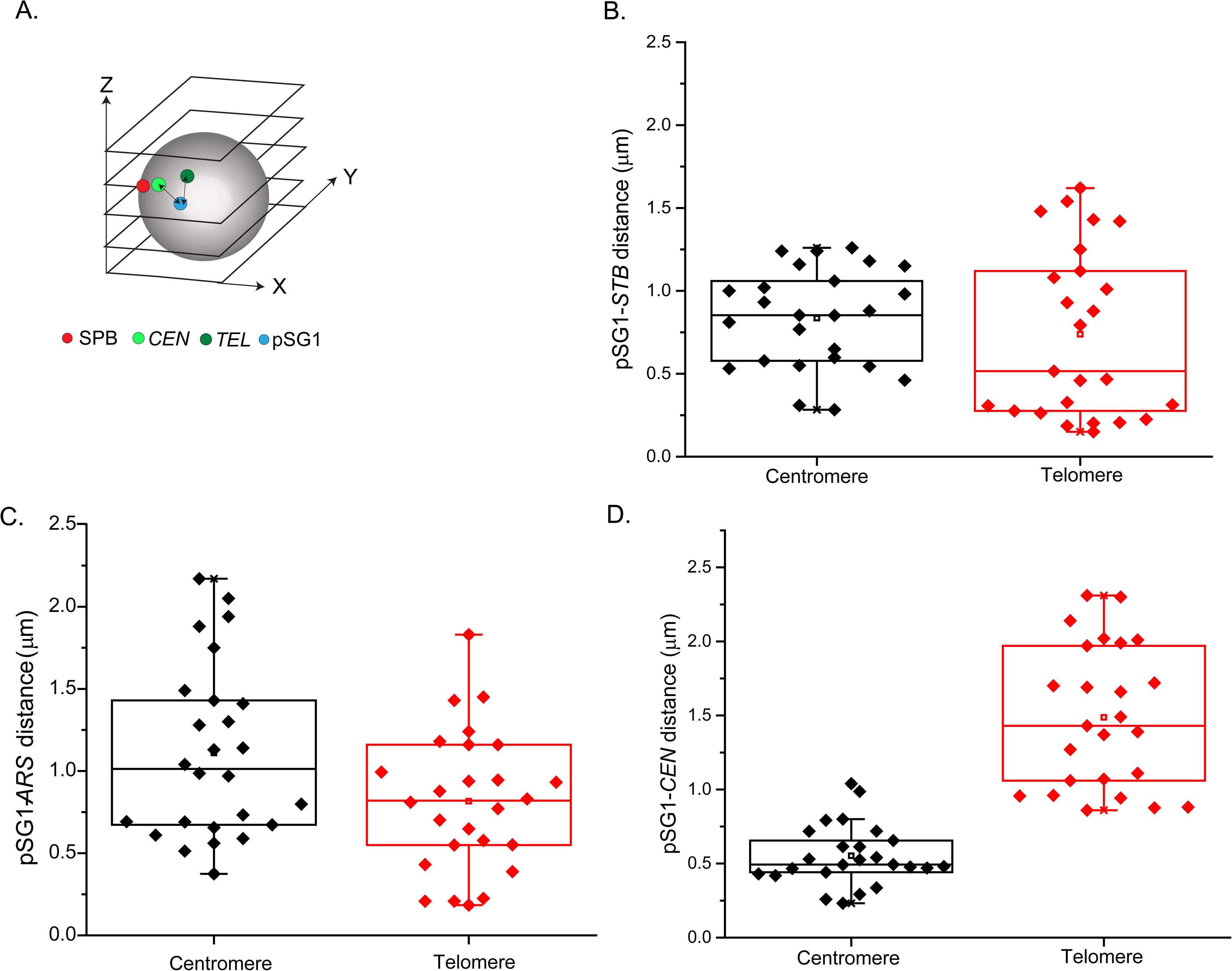
Relative distances of pSG1-*STB* from *CEN V* and *TEL V* within the 3D nuclear space. **A**. The positioning of the centroids of the fluorescent foci by Z-series sectioning of the nucleus is schematically shown. Images were captured from the [Cir^0^] experimental strain grown in glucose (pSG1-*CEN*) or galactose (pSG1-*ARS*) or the isogenic [Cir^+^] strain grown in galactose (pSG1-*STB*). Image analysis was performed using Imaris ‘Slice’ tool (see Materials and Methods for details). **B**-**D**. Plasmid distances from *CEN V* and *TEL V* are shown as dot plots.

**Figure 2- figure supplement 3.**
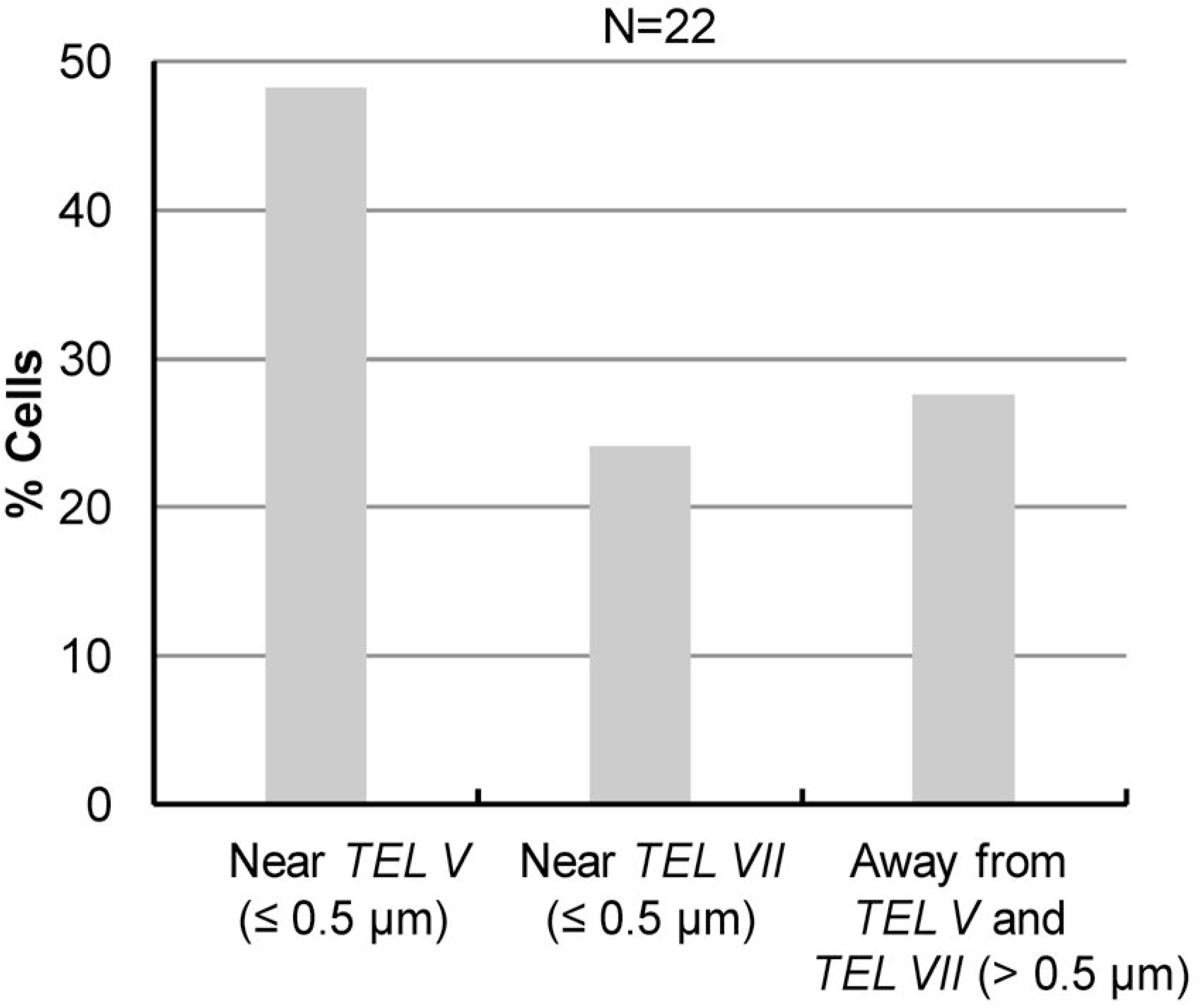
Simultaneous localization of pSG1*-STB* with respect to *TEL V* and *TEL VII*. The analysis was similar to that described in the legend to Figure 2. The pSG1-*STB* plasmid ([Cir^+^]; galactose; [LacO]_256_-[cyan-LacI]) was localized with respect to *TEL V* and *TEL VII* in fixed G1/S cells. The *TEL V* and *TEL VII* tagged by [TetO]_n_-[TetR-GFP] were distinguished by the difference in the brightness of their fluorescence (*TEL V* >> *TEL VII*). ‘N’ represents total number of analyzed cells.

**Figure 4- figure supplement 1.**
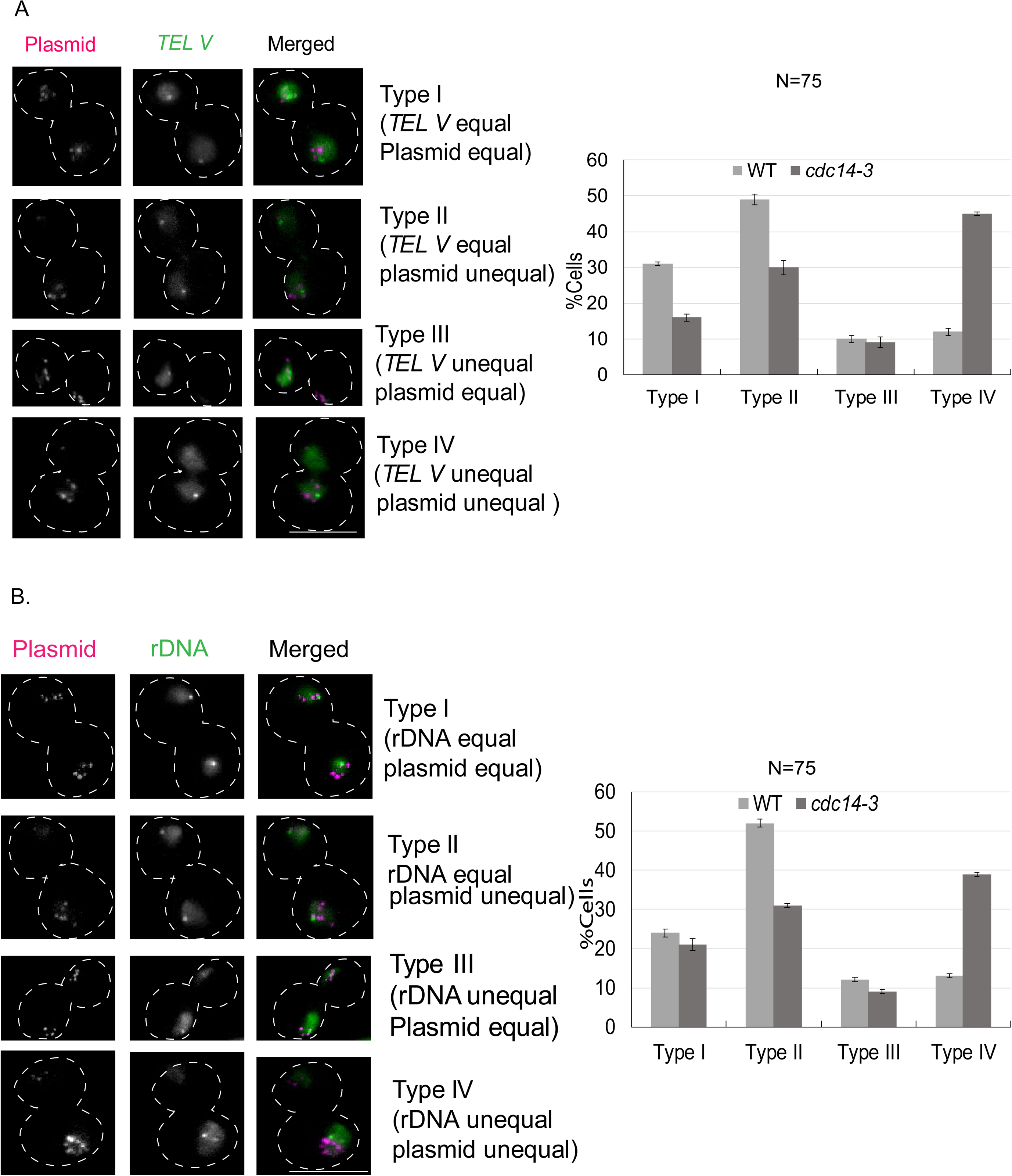
*ARS*-plasmid segregation when missegregation of telomeres or rDNA is induced by Cdc14 inactivation. **A**, **B**. The experimental protocols were similar to those described under Figure 4, except that the reporter plasmid employed was pSV6-*ARS*. Plasmid segregation was assayed in conjunction with that of *TEL V* (**A**) or of rDNA (**B**). The images of cell types (I-IV) and histogram plots of their quantitative analysis are arranged as in Figure 4. Bar, 5 µm. ‘N’ represents total number of cells analyzed. Error bars denote the standard deviation from the mean values acquired from three independent experiments.

**Figure 4- figure supplement 2.**
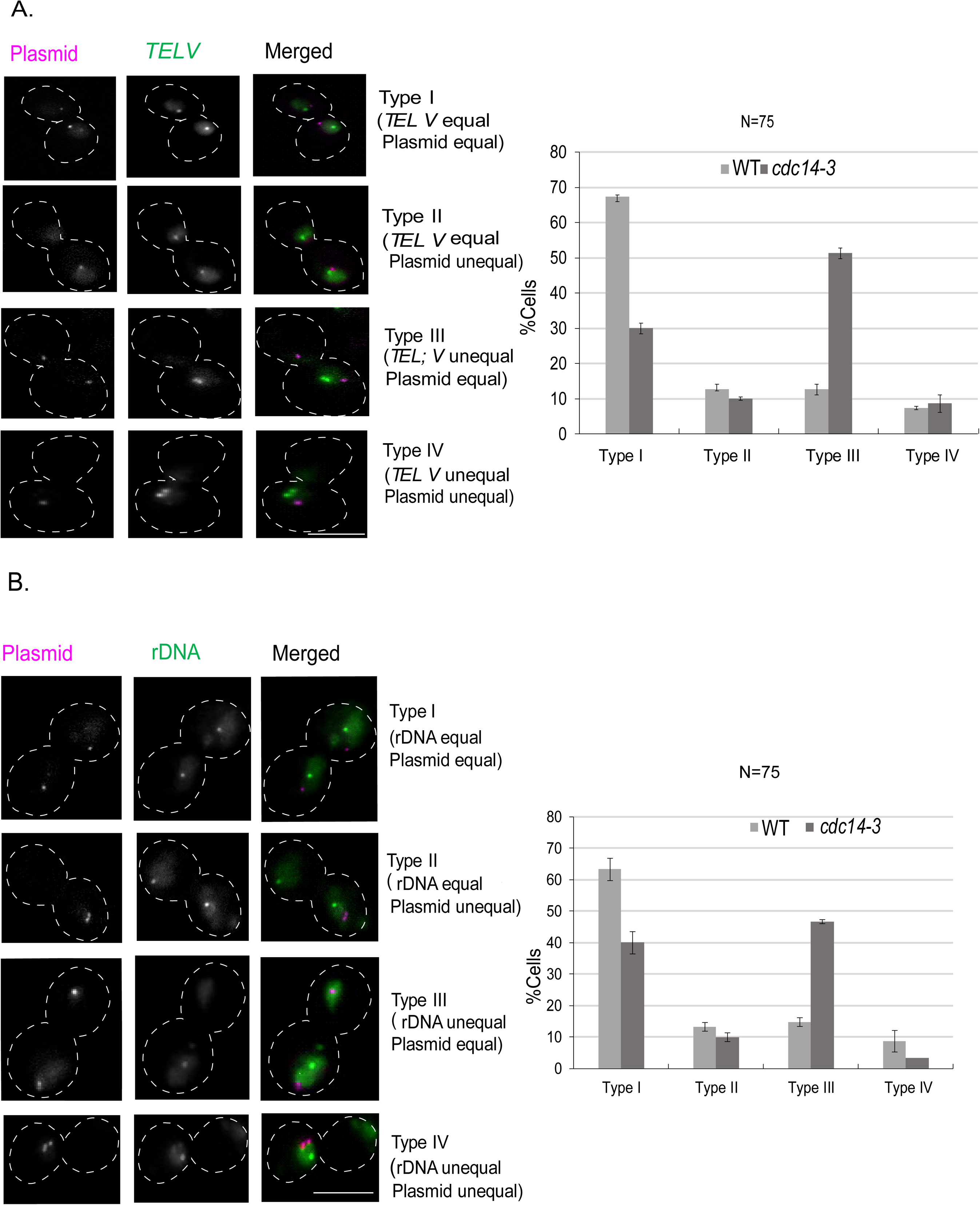
*CEN*-plasmid segregation under Cdc14 inactivation to induce missegregation of telomeres and rDNA. **A**, **B**. The assays in the wild type and *cdc-14-3* strains (otherwise isogenic) were carried out as described in the legend for Figure 4 (also Figure 4- figure supplement 1) with pSV7-*CEN* as the reporter plasmid. Bar, 5 μm. ‘N’ represents total number of cells analyzed. Error bars denote the standard deviation from the mean values acquired from three independent experiments.

**Figure 4- figure supplement 3.**
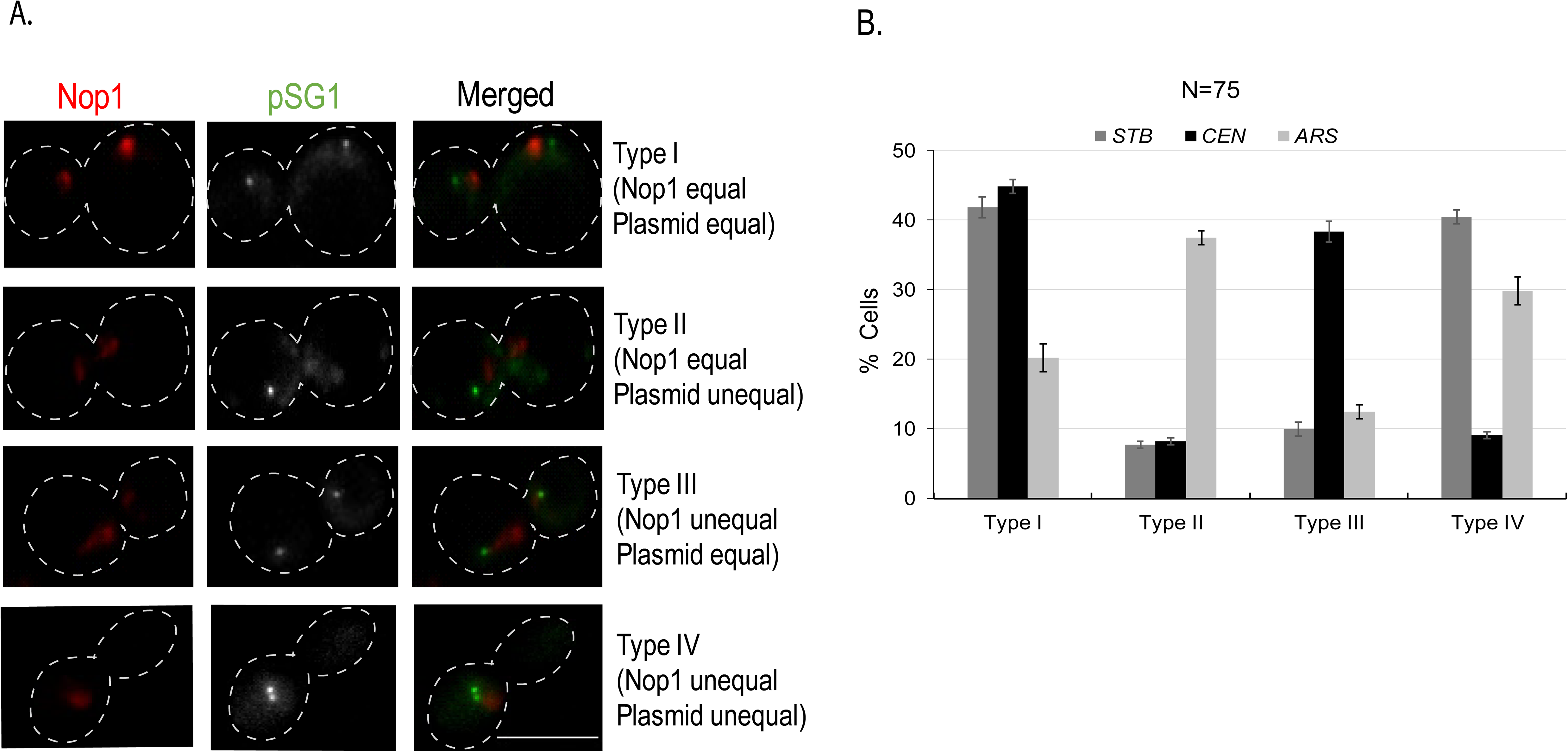
Segregation of single copy reporter plasmids (pSG1-*STB*, pSG1-*CEN*, and pSG1-*ARS*) with respect to Nop1 under Cdc14 inactivation. The partitioning status of the pSG1-reporter plasmid was manipulated as indicated in Figure 1- figure supplement 1 to obtain pSG1-*STB*, pSG1-*CEN* and pSG1-*ARS*. Segregation assays were performed at the non-permissive temperature (33°C). Segregation types (I-IV; images at the left) are quantitated in histogram plots at the right. Bar, 5μm. ‘N’ represents total number of cells analyzed. Error bars denote the standard deviation from the mean values acquired from three independent experiments. The correlation with Nop1 in segregation is high in the case of pSG1-*STB* (Type 1 + Type IV >> Type II + Type III), which is not the case for pSG1-*ARS* or pSG1-*CEN*. Equal segregation is high for pSG1-*CEN* (Type I + Type III >> Type II + Type IV); the opposite is true for pSG1-*ARS* (Type II + Type IV >> Type I + Type III).

**Figure 7- figure supplement 1.**
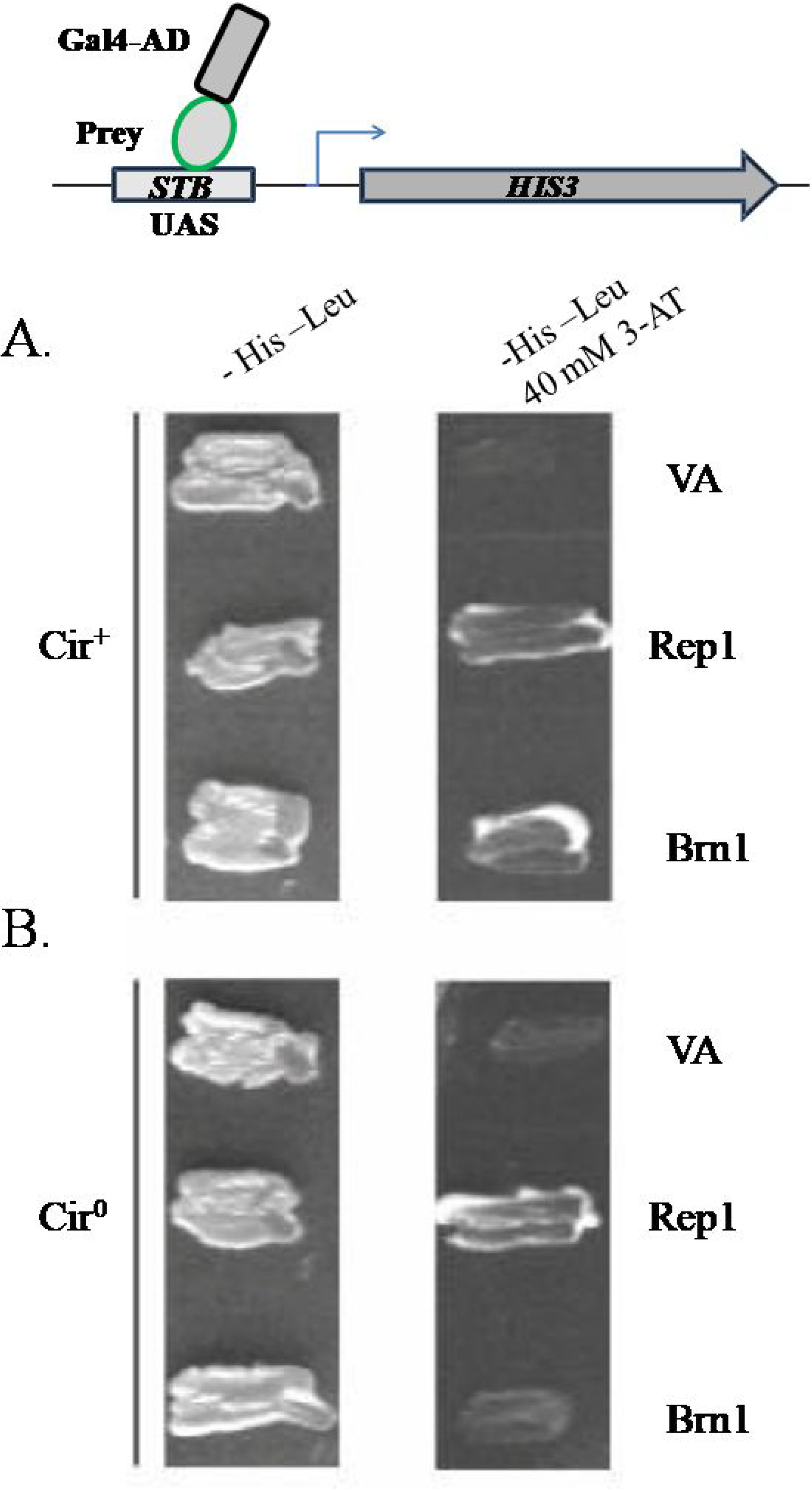
In a monohybrid assay, Brn1 interacts with the *STB* locus. The reporter construct for the monohybrid, schematically illustrated at the top, includes the *STB* locus as an artificial UAS (Upstream activating sequence) of the *HIS3* gene expressed from its native promoter. A test protein was expressed as a fusion to the *GAL4* activation domain under the control of the constitutive *ADH* promoter from a 2-micron derived plasmid vector (carrying the *LEU2* marker). The left and right columns show growth on plates lacking leucine and histidine in the absence and presence of 40 mM 3-AT (3-amino 1,2,4-triazole). The assays were performed in isogenic strains containing the 2-micron plasmid ([Cir^+^]) (**A**) or lacking it ([Cir^0^] (**B**). The latter strain lacks Rep1 and Rep2 expressed from the endogenous plasmid. Rep1, which is known to interact with *STB* (Sengupta et al., 2001) served as a positive control. The negative control is labeled ‘VA’, for vector alone.

**Figure 7- figure supplement 2.**
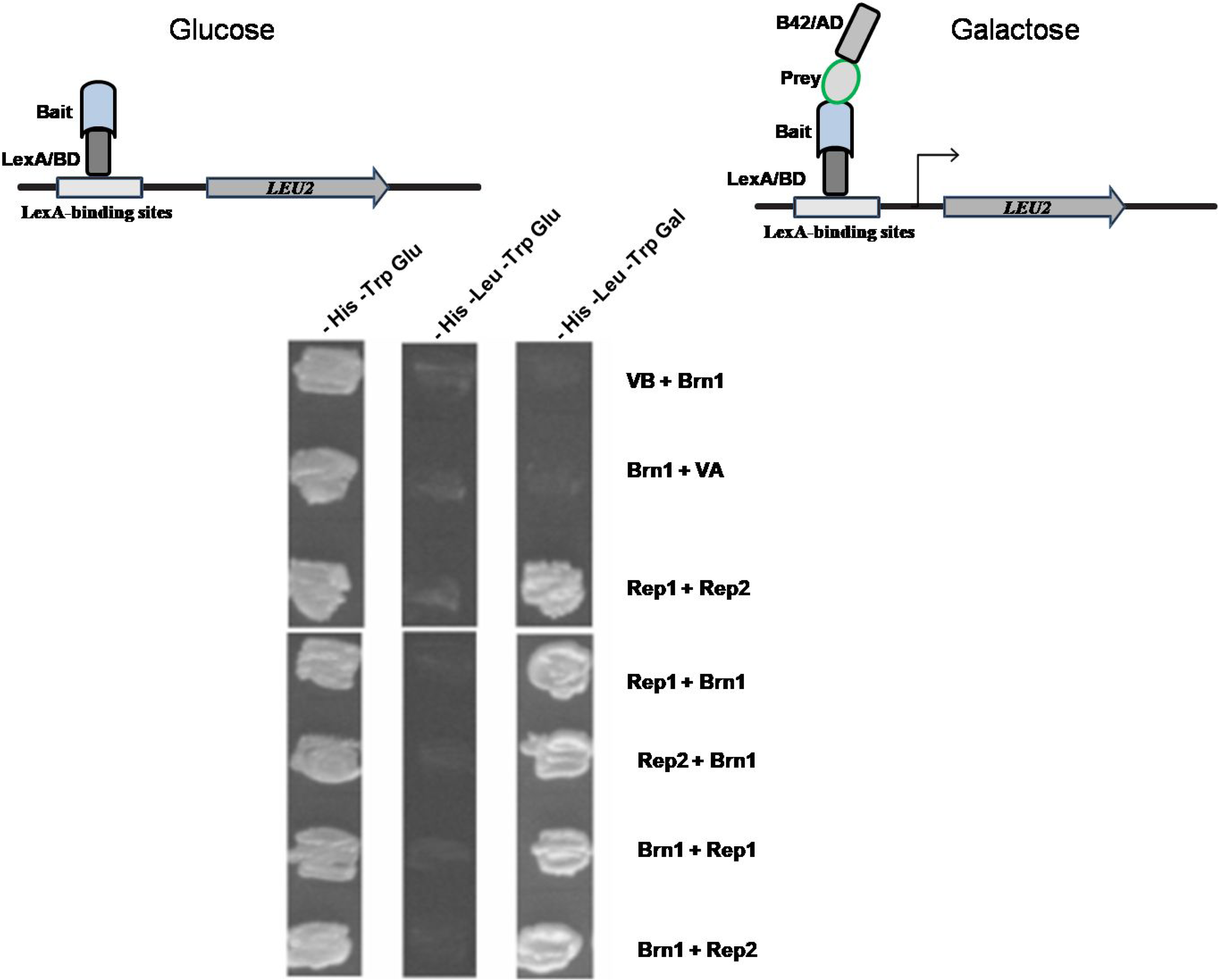
Brn1 interacts with Rep1 and Rep2 in dihybrid assays performed in a [Cir^+^] strain. The dihybrid system based on the *LEU2* reporter (Finely and Brent, 1997) is schematically diagrammed at the top. The ‘bait’ and ‘prey’ proteins were expressed as hybrid proteins fused to the LexA repressor and B42 activation domain, respectively. These *CEN*-based expression vectors were designed for inducible expression from the *GAL* promoter, and harbored *HIS3* (VB; bait expression vector) or *TRP1* (VA; prey expression vector) as selectable markers. For a given protein pair, the protein to the left of the plus sign was the bait; the one to the right was the potential prey. The Rep1 (bait)-Rep2 (prey) combination served as the positive control.

## References

A. Rizvi, S. M., Prajapati, H. K., Nag, P., & Ghosh, S. K. (2017). The 2-μm plasmid encoded protein Raf1 regulates both stability and copy number of the plasmid by blocking the formation of the Rep1–Rep2 repressor complex. Nucleic Acids Res, 45(12), 7167–7179. https://doi.org/10.1093/nar/gkx316

Alexandru, G., Uhlmann, F., Mechtler, K., Poupart, M.-A., & Nasmyth, K. (2001). Phosphorylation of the Cohesin Subunit Scc1 by Polo/Cdc5 Kinase Regulates Sister Chromatid Separation in Yeast. Cell, 105(4), 459–472. https://doi.org/10.1016/S0092-8674(01)00362-2

Autiero, M., Camarca, A., Ciullo, M., Debily, M. A., El Marhomy, S., Pasquinelli, R., Capasso, I., D’Aiuto, G., Anzisi, A. M., & Piatier-Tonneau, D. (2002). Intragenic amplification and formation of extrachromosomal small circular DNA molecules from the PIP gene on chromosome 7 in primary breast carcinomas. Int J Cancer, 99(3), 370–377.

Aydin, I., & Schelhaas, M. (2016). Viral Genome Tethering to Host Cell Chromatin: Cause and Consequences. Traffic, 17(4), 327–340. https://doi.org/10.1111/tra.12378

Biggins, S. (2013). The Composition, Functions, and Regulation of the Budding Yeast Kinetochore. Genetics, 194(4), 817. https://doi.org/10.1534/genetics.112.145276

Brachmann, C. B., Davies, A., Cost, G. J., Caputo, E., Li, J., Hieter, P., & Boeke, J. D. (1998). Designer deletion strains derived from Saccharomyces cerevisiae S288C: a useful set of strains and plasmids for PCR-mediated gene disruption and other applications. Yeast, 14(2), 115–132. https://doi.org/10.1002/(SICI)1097-0061(19980130)14:2<115::AID-YEA204>3.0.CO;2-2

Briggs, S. D., Bryk, M., Strahl, B. D., Cheung, W. L., Davie, J. K., Dent, S. Y., Winston, F., & Allis, C. D. (2001). Histone H3 lysine 4 methylation is mediated by Set1 and required for cell growth and rDNA silencing in Saccharomyces cerevisiae. Genes Dev, 15(24), 3286–3295. https://doi.org/10.1101/gad.940201

Broach, J. R. (1982). The yeast plasmid 2 mu circle. Cell, 28(2), 203–204. https://doi.org/10.1016/0092-8674(82)90337-3

Broach, J. R., & Volkert, F. C. (1991). 6 Circular DNA Plasmids of Yeasts. Cold Spring Harbor Monograph Archive, 21, 297–331. http://dx.doi.org/10.1101/0.297-331

Bystricky, K., Laroche, T., van Houwe, G., Blaszczyk, M., & Gasser, S. M. (2005). Chromosome looping in yeast: telomere pairing and coordinated movement reflect anchoring efficiency and territorial organization. J Cell Biol, 168(3), 375–387. https://doi.org/10.1083/jcb.200409091

Charbin, A., Bouchoux, C., & Uhlmann, F. (2014). Condensin aids sister chromatid decatenation by topoisomerase II. Nucleic Acids Res, 42(1), 340–348. https://doi.org/10.1093/nar/gkt882

Chen, X. L., Reindle, A., & Johnson, E. S. (2005). Misregulation of 2μm circle copy number in a SUMO pathway mutant. Mol Cell Biol, 25(10), 4311–4320. https://dx.doi.org/10.1128/2FMCB.25.10.4311-4320.2005

Chu, S. (1998). The transcriptional program of sporulation in budding yeast (vol 282, pg 699, 1998). Science, 282(5393), 1421–1421. https://doi.org/10.1126/science.282.5389.699

Chua, P. R., & Roeder, G. S. (1997). Tam1, a telomere-associated meiotic protein, functions in chromosome synapsis and crossover interference. Genes Dev, 11(14), 1786–1800. https://doi.org/10.1101/gad.11.14.1786

Clemente-Blanco, A., Sen, N., Mayan-Santos, M., Sacristan, M. P., Graham, B., Jarmuz, A., Giess, A., Webb, E., Game, L., Eick, D., Bueno, A., Merkenschlager, M., & Aragon, L. (2011). Cdc14 phosphatase promotes segregation of telomeres through repression of RNA polymerase II transcription. Nat Cell Biol, 13(12), 1450–1456. https://doi.org/10.1038/ncb2365

Cohen, S., Regev, A., & Lavi, S. (1997). Small polydispersed circular DNA (spcDNA) in human cells: association with genomic instability. Oncogene, 14(8), 977–985. https://doi.org/10.1038/sj.onc.1200917

Cohen, S., & Segal, D. (2009). Extrachromosomal circular DNA in eukaryotes: possible involvement in the plasticity of tandem repeats. Cytogenet Genome Res, 124(3-4), 327–338. https://doi.org/10.1159/000218136

Conrad, M. N., Dominguez, A. M., & Dresser, M. E. (1997). Ndj1p, a meiotic telomere protein required for normal chromosome synapsis and segregation in yeast. Science, 276(5316), 1252–1255. https://doi.org/10.1126/science.276.5316.1252

Coursey, T. L., & McBride, A. A. (2019). Hitchhiking of Viral Genomes on Cellular Chromosomes. Annu Rev Virol, 6(1), 275–296. https://doi.org/10.1146/annurev-virology-092818-015716

Cui, H., Ghosh, S. K., & Jayaram, M. (2009). The selfish yeast plasmid uses the nuclear motor Kip1p but not Cin8p for its localization and equal segregation. J Cell Biol, 185(2), 251–264. https://doi.org/10.1083/jcb.200810130

D’Amours, D., Stegmeier, F., & Amon, A. (2004). Cdc14 and condensin control the dissolution of cohesin-independent chromosome linkages at repeated DNA. Cell, 117(4), 455–469. https://doi.org/10.1016/s0092-8674(04)00413-1

DiNardo, S., Voelkel, K., & Sternglanz, R. (1984). DNA topoisomerase II mutant of Saccharomyces cerevisiae: topoisomerase II is required for segregation of daughter molecules at the termination of DNA replication.PNAS, 81(9), 2616–2620. https://dx.doi.org/10.1073/pnas.81.9.2616

Fennell, A., Fernández-Álvarez, A., Tomita, K., & Cooper, J. P. (2015). Telomeres and centromeres have interchangeable roles in promoting meiotic spindle formation. J Cell Biol, 208(4), 415–428. https://dx.doi.org/10.1083/jcb.201409058

Finley, R. L., Thomas, B. J., Zipursky, S. L., & Brent, R. (1996). Isolation of Drosophila cyclin D, a protein expressed in the morphogenetic furrow before entry into S phase.PNAS, 93(7), 3011–3015. https://dx.doi.org/10.1073/2Fpnas.93.7.3011

Frappier, L. (2004). Viral plasmids in mammalian cells. In Plasmid Biology (pp. 325–340). American Society of Microbiology. Chapter DOI: 10.1128/9781555817732.ch15

Futcher, A. B. (1986). Copy number amplification of the 2 micron circle plasmid of Saccharomyces cerevisiae. J Theor Biol, 119(2), 197–204. https://doi.org/10.1016/s0022-5193(86)80074-1

Futcher, A. B., & Cox, B. S. (1983). Maintenance of the 2 microns circle plasmid in populations of Saccharomyces cerevisiae. J Bacteriol, 154(2), 612–622. http://www.ncbi.nlm.nih.gov/pubmed/6341357

Ghosh, S. K., Hajra, S., & Jayaram, M. (2007). Faithful segregation of the multicopy yeast plasmid through cohesin-mediated recognition of sisters.PNAS, 104(32), 13034–13039. https://doi.org/10.1073/pnas.0702996104

Ghosh, S. K., Hajra, S., Paek, A., & Jayaram, M. (2006). Mechanisms for chromosome and plasmid segregation. Annu Rev Biochem, 75, 211–241. https://doi.org/10.1146/annurev.biochem.75.101304.124037

Ghosh, S. K., Huang, C. C., Hajra, S., & Jayaram, M. (2010). Yeast cohesin complex embraces 2 micron plasmid sisters in a tri-linked catenane complex. Nucleic Acids Res, 38(2), 570–584. https://doi.org/10.1093/nar/gkp993

Gotta, M., Laroche, T., Formenton, A., Maillet, L., Scherthan, H., & Gasser, S. M. (1996). The clustering of telomeres and colocalization with Rap1, Sir3, and Sir4 proteins in wild-type Saccharomyces cerevisiae. J Cell Biol, 134(6), 1349–1363. https://doi.org/10.1083/jcb.134.6.1349

Griffiths, R., & Whitehouse, A. (2007). Herpesvirus saimiri episomal persistence is maintained via interaction between open reading frame 73 and the cellular chromosome-associated protein MeCP2. J Virol, 81(8), 4021–4032. https://doi.org/10.1128/jvi.02171-06

Grunstein, M., & Gasser, S. M. (2013). Epigenetics in Saccharomyces cerevisiae. Cold Spring Harb Perspect Biol, 5(7), a017491. https://doi.org/10.1101/cshperspect.a017491

Guacci, V., Hogan, E., & Koshland, D. (1997). Centromere position in budding yeast: evidence for anaphase A. Mol Biol Cell, 8(6), 957–972. https://doi.org/10.1091/mbc.8.6.957

Haeusler, R. A., Pratt-Hyatt, M., Good, P. D., Gipson, T. A., & Engelke, D. R. (2008). Clustering of yeast tRNA genes is mediated by specific association of condensin with tRNA gene transcription complexes. Genes Dev, 22(16), 2204–2214. https://doi.org/10.1101/gad.1675908

Hajra, S., Ghosh, S. K., & Jayaram, M. (2006). The centromere-specific histone variant Cse4p (CENP-A) is essential for functional chromatin architecture at the yeast 2-microm circle partitioning locus and promotes equal plasmid segregation. J Cell Biol, 174(6), 779–790. https://doi.org/10.1083/jcb.200603042

Hayes, F., & Barilla, D. (2006). The bacterial segrosome: a dynamic nucleoprotein machine for DNA trafficking and segregation [Review Article]. Nat Rev Microbiol, 4(2), 133–143. https://doi.org/10.1038/nrmicro1342

Heun, P., Laroche, T., Raghuraman, M. K., & Gasser, S. M. (2001a). The positioning and dynamics of origins of replication in the budding yeast nucleus. J Cell Biol, 152(2), 385–400. http://www.ncbi.nlm.nih.gov/pubmed/11266454

Heun, P., Laroche, T., Shimada, K., Furrer, P., & Gasser, S. M. (2001b). Chromosome dynamics in the yeast interphase nucleus. Science, 294(5549), 2181–2186. https://doi.org/10.1126/science.1065366

Hill, A., & Bloom, K. (1989). Acquisition and processing of a conditional dicentric chromosome in Saccharomyces cerevisiae. Mol Cell Biol, 9(3), 1368–1370. https://doi.org/10.1128/mcb.9.3.1368

Huang, C. C., Chang, K. M., Cui, H., & Jayaram, M. (2011). Histone H3-variant Cse4-induced positive DNA supercoiling in the yeast plasmid has implications for a plasmid origin of a chromosome centromere.PNAS, 108(33), 13671–13676. https://doi.org/10.1073/pnas.1101944108

Huang, Y., Xu, L., Sun, Y., & Nabel, G. J. (2002). The assembly of Ebola virus nucleocapsid requires virion-associated proteins 35 and 24 and posttranslational modification of nucleoprotein. Mol Cell, 10(2), 307–316. https://doi.org/10.1016/s1097-2765(02)00588-9

Hughes, J. E., & Welker, D. L. (1999). Nuclear plasmids of Dictyostelium. Genet Eng (N Y), 21, 1–14. https://doi.org/10.1007/978-1-4615-4707-5_1

Janke, C., Magiera, M. M., Rathfelder, N., Taxis, C., Reber, S., Maekawa, H., Moreno-Borchart, A., Doenges, G., Schwob, E., & Schiebel, E. (2004). A versatile toolbox for PCR-based tagging of yeast genes: new fluorescent proteins, more markers and promoter substitution cassettes. Yeast, 21(11), 947–962. https://doi.org/10.1002/yea.1142

Jin, Q.-W., Fuchs, J., & Loidl, J. (2000). Centromere clustering is a major determinant of yeast interphase nuclear organization. J Cell Sci, 113(11), 1903–1912. http://www.ncbi.nlm.nih.gov/pubmed/10806101

Jin, Q.-w., Trelles-Sticken, E., Scherthan, H., & Loidl, J. (1998). Yeast nuclei display prominent centromere clustering that is reduced in nondividing cells and in meiotic prophase. J Cell Biol, 141(1), 21–29. https://doi.org/10.1083/jcb.141.1.21

Kanda, T., Kamiya, M., Maruo, S., Iwakiri, D., & Takada, K. (2007). Symmetrical localization of extrachromosomally replicating viral genomes on sister chromatids. J Cell Sci, 120(Pt 9), 1529–1539. https://doi.org/10.1242/jcs.03434

Kelley-Clarke, B., Ballestas, M. E., Komatsu, T., & Kaye, K. M. (2007). Kaposi’s sarcoma herpesvirus C-terminal LANA concentrates at pericentromeric and peri-telomeric regions of a subset of mitotic chromosomes. Virology, 357(2), 149–157. https://doi.org/10.1016/j.virol.2006.07.052

Khmelinskii, A., & Schiebel, E. (2008). Assembling the spindle midzone in the right place at the right time. Cell Cycle, 7(3), 283–286. https://doi.org/10.4161/cc.7.3.5349

Krithivas, A., Fujimuro, M., Weidner, M., Young, D. B., & Hayward, S. D. (2002). Protein interactions targeting the latency-associated nuclear antigen of Kaposi’s sarcoma-associated herpesvirus to cell chromosomes. J Virol, 76(22), 11596–11604. https://dx.doi.org/10.1128/JVI.76.22.11596-11604.2002

Kruitwagen, T., Chymkowitch, P., Denoth-Lippuner, A., Enserink, J., & Barral, Y. (2018). Centromeres License the Mitotic Condensation of Yeast Chromosome Arms. Cell, 175(3), 780–795 e715. https://doi.org/10.1016/j.cell.2018.09.012

Kueng, S., Oppikofer, M., & Gasser, S. M. (2013). SIR proteins and the assembly of silent chromatin in budding yeast. Annu Rev Genet, 47, 275–306. https://doi.org/10.1146/annurev-genet-021313-173730

Kupiec, M. (2014). Biology of telomeres: lessons from budding yeast. FEMS Microbiol Rev, 38(2), 144–171. https://doi.org/10.1111/1574-6976.12054

Lavoie, B. D., Hogan, E., & Koshland, D. (2004). In vivo requirements for rDNA chromosome condensation reveal two cell-cycle-regulated pathways for mitotic chromosome folding. Genes Dev, 18(1), 76–87. https://doi.org/10.1101/gad.1150404

Lefrancois, P., Auerbach, R. K., Yellman, C. M., Roeder, G. S., & Snyder, M. (2013). Centromere-like regions in the budding yeast genome. PLoS Genet, 9(1), e1003209. https://doi.org/10.1371/journal.pgen.1003209

Liu, Y. T., Chang, K. M., Ma, C. H., & Jayaram, M. (2016). Replication-dependent and independent mechanisms for the chromosome-coupled persistence of a selfish genome. Nucleic Acids Res, 44(17), 8302–8323. https://doi.org/10.1093/nar/gkw694

Liu, Y. T., Ma, C. H., & Jayaram, M. (2013). Co-segregation of yeast plasmid sisters under monopolin-directed mitosis suggests association of plasmid sisters with sister chromatids. Nucleic Acids Res, 41(7), 4144–4158. https://doi.org/10.1093/nar/gkt096

Liu, Y. T., Sau, S., Ma, C. H., Kachroo, A. H., Rowley, P. A., Chang, K. M., Fan, H. F., & Jayaram, M. (2014). The partitioning and copy number control systems of the selfish yeast plasmid: an optimized molecular design for stable persistence in host cells. Microbiol Spectr, 2(5). https://doi.org/10.1128/microbiolspec.PLAS-0003-2013

Ma, C.-H., Cui, H., Hajra, S., Rowley, P. A., Fekete, C., Sarkeshik, A., Ghosh, S. K., Yates III, J. R., & Jayaram, M. (2012). Temporal sequence and cell cycle cues in the assembly of host factors at the yeast 2 micron plasmid partitioning locus. Nucleic Acids Res, 41(4), 2340–2353. http://doi.org/10.1093/nar/gks1338

Ma, C.-H., Su, B.-Y., Maciaszek, A., Fan, H.-F., Guga, P., & Jayaram, M. (2019). A Flp-SUMO hybrid recombinase reveals multi-layered copy number control of a selfish DNA element through post-translational modification. PLoS Genet, 15(6), e1008193. https://doi.org/10.1371/journal.pgen.1008193

Ma, C. H., Cui, H., Hajra, S., Rowley, P. A., Fekete, C., Sarkeshik, A., Ghosh, S. K., Yates, J. R., 3rd, & Jayaram, M. (2013). Temporal sequence and cell cycle cues in the assembly of host factors at the yeast 2 micron plasmid partitioning locus. Nucleic Acids Res, 41(4), 2340–2353. https://doi.org/10.1093/nar/gks1338

Machin, F., Quevedo, O., Ramos-Perez, C., & Garcia-Luis, J. (2016). Cdc14 phosphatase: warning, no delay allowed for chromosome segregation! Curr Genet, 62(1), 7–13. https://doi.org/10.1007/s00294-015-0502-1

Malik, H. S., & Henikoff, S. (2009). Major evolutionary transitions in centromere complexity. Cell, 138(6), 1067–1082. https://doi.org/10.1016/j.cell.2009.08.036

Marston, A. L., Lee, B. H., & Amon, A. (2003). The Cdc14 phosphatase and the FEAR network control meiotic spindle disassembly and chromosome segregation. Dev Cell, 4(5), 711–726. https://doi.org/10.1016/S1534-5807(03)00130-8

McBride, A. A. (2008). Replication and partitioning of papillomavirus genomes. Adv Virus Res, 72, 155–205. https://doi.org/10.1016/S0065-3527(08)00404-1

Mead, D. J., Gardner, D. C., & Oliver, S. G. (1986). The yeast 2 μ plasmid: strategies for the survival of a selfish DNA. Mol Gen Genet 205(3), 417–421. https://doi.org/10.1007/BF00338076

Mehta, S., Yang, X. M., Chan, C. S., Dobson, M. J., Jayaram, M., & Velmurugan, S. (2002). The 2 micron plasmid purloins the yeast cohesin complex: a mechanism for coupling plasmid partitioning and chromosome segregation? J Cell Biol, 158(4), 625–637. https://doi.org/10.1083/jcb.200204136

Mehta, S., Yang, X. M., Jayaram, M., & Velmurugan, S. (2005). A novel role for the mitotic spindle during DNA segregation in yeast: promoting 2 microm plasmid-cohesin association. Mol Cell Biol, 25(10), 4283–4298. https://doi.org/10.1128/MCB.25.10.4283-4298.2005

Michaelis, C., Ciosk, R., & Nasmyth, K. (1997). Cohesins: chromosomal proteins that prevent premature separation of sister chromatids. Cell, 91(1), 35–45. https://doi.org/10.1016/s0092-8674(01)80007-6

Mittal P., Ghule K., Trakroo D., Prajapati H.K., & Ghosh, S.K. (2020). Meiosis-specific functions of kinesin motors in cohesin removal and maintenance of chromosome integrity in budding yeast. Mol Cell Biol; 10.1128/MCB.00386-19. https://doi.org/10.1128/MCB.00386-19

Møller, H. D., Parsons, L., Jørgensen, T. S., Botstein, D., & Regenberg, B. (2015). Extrachromosomal circular DNA is common in yeast. PNAS, 112(24), E3114–E3122. https://doi.org/10.1073/pnas.1508825112

Moquin, S. A., Thomas, S., Whalen, S., Warburton, A., Fernandez, S. G., McBride, A. A., Pollard, K. S., & Miranda, J. J. L. (2017). The Epstein-Barr virus episome maneuvers between nuclear chromatin compartments during reactivation. J Virol. https://doi.org/10.1128/jvi.01413-17

Murray, J. A., & Cesareni, G. (1986). Functional analysis of the yeast plasmid partition locus STB. The EMBO journal, 5(12), 3391–3399. http://www.ncbi.nlm.nih.gov/pubmed/16453734

Murray, J. A., Scarpa, M., Rossi, N., & Cesareni, G. (1987). Antagonistic controls regulate copy number of the yeast 2 mu plasmid. The EMBO journal, 6(13), 4205–4212. http://www.ncbi.nlm.nih.gov/pubmed/2832156

Mythreye, K., & Bloom, K. S. (2003). Differential kinetochore protein requirements for establishment versus propagation of centromere activity in Saccharomyces cerevisiae. J Cell Biol, 160(6), 833–843. https://doi.org/10.1083/jcb.200211116

Nanbo, A., Sugden, A., & Sugden, B. (2007). The coupling of synthesis and partitioning of EBV’s plasmid replicon is revealed in live cells. The EMBO journal, 26(19), 4252–4262. https://doi.org/10.1038/sj.emboj.7601853

Ohkuni, K., & Kitagawa, K. (2012). Role of transcription at centromeres in budding yeast. Transcription, 3(4), 193–197. https://doi.org/10.4161/trns.20884

Oliveira, J. G., Colf, L. A., & McBride, A. A. (2006). Variations in the association of papillomavirus E2 proteins with mitotic chromosomes.PNAS, 103(4), 1047–1052. https://doi.org/10.1073/pnas.0507624103

Ouspenski, I. I., Cabello, O. A., & Brinkley, B. R. (2000). Chromosome condensation factor Brn1p is required for chromatid separation in mitosis. Mol Biol Cell, 11(4), 1305–1313. https://doi.org/10.1091/mbc.11.4.1305

Petes, T. D., & Williamson, D. H. (1994). A novel structural form of the 2 micron plasmid of the yeast Saccharomyces cerevisiae. Yeast, 10(10), 1341–1345. https://doi.org/10.1002/yea.320101011

Poddar, A., Reed, S. C., McPhillips, M. G., Spindler, J. E., & McBride, A. A. (2009). The human papillomavirus type 8 E2 tethering protein targets the ribosomal DNA loci of host mitotic chromosomes. J Virol, 83(2), 640–650. https://doi.org/10.1128/jvi.01936-08

Prajapati, H. K., Agarwal, M., Mittal, P., & Ghosh, S. K. (2018). Evidence of Zip1 Promoting Sister Kinetochore Mono-orientation During Meiosis in Budding Yeast. G3 (Bethesda, Md.), 8(11), 3691–3701. https://doi.org/10.1534/g3.118.200469

Prajapati, H. K., Rizvi, S. M. A., Rathore, I., & Ghosh, S. K. (2017). Microtubule-associated proteins, Bik1 and Bim1, are required for faithful partitioning of the endogenous 2 micron plasmids in budding yeast. Mol Microbiol, 103(6), 1046–1064. https://doi.org/10.1111/mmi.13608

Primig, M., Williams, R. M., Winzeler, E. A., Tevzadze, G. G., Conway, A. R., Hwang, S. Y., Davis, R. W., & Esposito, R. E. (2000). The core meiotic transcriptome in budding yeasts. Nat Genet, 26(4), 415–423. https://doi.org/10.1038/82539

Rabitsch, K. P., Tóth, A., Gálová, M., Schleiffer, A., Schaffner, G., Aigner, E., Rupp, C., Penkner, A. M., Moreno-Borchart, A. C., & Primig, M. (2001). A screen for genes required for meiosis and spore formation based on whole-genome expression. Curr Biol, 11(13), 1001–1009. https://doi.org/10.1016/s0960-9822(01)00274-3

Reid, R. J., Sunjevaric, I., Voth, W. P., Ciccone, S., Du, W., Olsen, A. E., Stillman, D. J., & Rothstein, R. (2008). Chromosome-scale genetic mapping using a set of 16 conditionally stable Saccharomyces cerevisiae chromosomes. Genetics, 180(4), 1799–1808. https://dx.doi.org/10.1534/Fgenetics.108.087999

Reynolds, A. E., Murray, A. W., & Szostak, J. W. (1987). Roles of the 2 microns gene products in stable maintenance of the 2 microns plasmid of Saccharomyces cerevisiae.Mol Cell Biol, 7(10), 3566–3573. https://doi.org/10.1128/mcb.7.10.3566

Rizvi, S. M. A., Prajapati, H. K., & Ghosh, S. K. (2018). The 2 micron plasmid: a selfish genetic element with an optimized survival strategy within Saccharomyces cerevisiae. Curr Genet, 64(1), 25–42. https://doi.org/10.1007/s00294-017-0719-2

Sau, S., Conrad, M. N., Lee, C. Y., Kaback, D. B., Dresser, M. E., & Jayaram, M. (2014). A selfish DNA element engages a meiosis-specific motor and telomeres for germ-line propagation. J Cell Biol, 205(5), 643–661. https://doi.org/10.1083/jcb.201312002

Sau, S., Ghosh, S. K., Liu, Y. T., Ma, C. H., & Jayaram, M. (2019). Hitchhiking on chromosomes: A persistence strategy shared by diverse selfish DNA elements. Plasmid, 102, 19–28. https://doi.org/10.1016/j.plasmid.2019.01.004

Sau, S., Liu, Y. T., Ma, C. H., & Jayaram, M. (2015). Stable persistence of the yeast plasmid by hitchhiking on chromosomes during vegetative and germ-line divisions of host cells. Mob Genet Elements, 5(2), 1–8. https://doi.org/10.1080/2159256x.2015.1031359

Scott-Drew, S., & Murray, J. (1998). Localisation and interaction of the protein components of the yeast 2 mu circle plasmid partitioning system suggest a mechanism for plasmid inheritance. J Cell Sci, 111(13), 1779–1789. http://www.ncbi.nlm.nih.gov/pubmed/9625741

Sekhar, V., Reed, S. C., & McBride, A. A. (2010). Interaction of the betapapillomavirus E2 tethering protein with mitotic chromosomes. J Virol, 84(1), 543–557. https://doi.org/10.1128/jvi.01908-09

Sharp, J. A., Krawitz, D. C., Gardner, K. A., Fox, C. A., & Kaufman, P. D. (2003). The budding yeast silencing protein Sir1 is a functional component of centromeric chromatin. Genes Dev, 17(19), 2356–2361. https://doi.org/10.1101/gad.1131103

Shibata, Y., Kumar, P., Layer, R., Willcox, S., Gagan, J. R., Griffith, J. D., & Dutta, A. (2012). Extrachromosomal microDNAs and chromosomal microdeletions in normal tissues. Science, 336(6077), 82–86. https://doi.org/10.1126/science.1213307

Som, T., Armstrong, K. A., Volkert, F. C., & Broach, J. R. (1988). Autoregulation of 2μm circle gene expression provides a model for maintenance of stable plasmid copy levels. Cell, 52(1), 27–37. https://doi.org/10.1016/0092-8674(88)90528-4

Stegmeier, F., & Amon, A. (2004). Closing mitosis: the functions of the Cdc14 phosphatase and its regulation. Annu Rev Genet, 38, 203–232. https://doi.org/10.1146/annurev.genet.38.072902.093051

Stephens, A. D., Haase, J., Vicci, L., Taylor, R. M., 2nd, & Bloom, K. (2011). Cohesin, condensin, and the intramolecular centromere loop together generate the mitotic chromatin spring. J Cell Biol, 193(7), 1167–1180. https://doi.org/10.1083/jcb.201103138

Straight, A. F., Belmont, A. S., Robinett, C. C., & Murray, A. W. (1996). GFP tagging of budding yeast chromosomes reveals that protein-protein interactions can mediate sister chromatid cohesion. Curr Biol, 6(12), 1599–1608. https://doi.org/10.1016/S0960-9822(02)70783-5

Sullivan, M., Higuchi, T., Katis, V. L., & Uhlmann, F. (2004). Cdc14 phosphatase induces rDNA condensation and resolves cohesin-independent cohesion during budding yeast anaphase. Cell, 117(4), 471–482. https://doi.org/10.1016/s0092-8674(04)00415-5

Taddei, A., & Gasser, S. M. (2012). Structure and function in the budding yeast nucleus. Genetics, 192(1), 107–129. https://doi.org/10.1534/genetics.112.140608

Takeo, S., Lake, C. M., Morais-de-Sá, E., Sunkel, C. E., & Hawley, R. S. (2011). Synaptonemal complex-dependent centromeric clustering and the initiation of synapsis in Drosophila oocytes. Curr Biol, 21(21), 1845–1851. https://doi.org/10.1016/j.cub.2011.09.044

Velmurugan, S., Yang, X.-M., Chan, C. S.-M., Dobson, M., & Jayaram, M. (2000). Partitioning of the 2-μm circle plasmid of Saccharomyces cerevisiae: functional coordination with chromosome segregation and plasmid-encoded Rep protein distribution. J Cell Biol, 149(3), 553–566. https://doi.org/10.1083/jcb.149.3.553

Verzijlbergen, K. F., Nerusheva, O. O., Kelly, D., Kerr, A., Clift, D., de Lima Alves, F., Rappsilber, J., & Marston, A. L. (2014). Shugoshin biases chromosomes for biorientation through condensin recruitment to the pericentromere. eLife, 3, e01374. https://doi.org/10.7554/eLife.01374

Villasante, A., Abad, J. P., & Méndez-Lago, M. (2007). Centromeres were derived from telomeres during the evolution of the eukaryotic chromosome.PNAS, 104(25), 10542–10547. https://dx.doi.org/10.1073/pnas.0703808104

Volkert, F. C., & Broach, J. R. (1986). Site-specific recombination promotes plasmid amplification in yeast. Cell, 46(4), 541–550. https://doi.org/10.1016/0092-8674(86)90879-2

Wang, B. D., Eyre, D., Basrai, M., Lichten, M., & Strunnikov, A. (2005). Condensin binding at distinct and specific chromosomal sites in the Saccharomyces cerevisiae genome. Mol Cell Biol, 25(16), 7216–7225. https://doi.org/10.1128/MCB.25.16.7216-7225.2005

Wong, M. C. V. L., Scott-Drew, S. R. S., Hayes, M. J., Howard, P. J., & Murray, J. A. H. (2002). RSC2, encoding a component of the RSC nucleosome remodeling complex, is essential for 2 microm plasmid maintenance in Saccharomyces cerevisiae. Mol Cell Biol, 22(12), 4218–4229. https://doi.org/10.1128/mcb.22.12.4218-4229.2002

Xiong, L., Chen, X. L., Silver, H. R., Ahmed, N. T., & Johnson, E. S. (2009). Deficient SUMO attachment to Flp recombinase leads to homologous recombination-dependent hyperamplification of the yeast 2 μm circle plasmid.Mol Biol Cell, 20(4), 1241–1251. https://dx.doi.org/10.1091/mbc.E08-06-0659

Yong-Gonzalez, V., Wang, B.-D., Butylin, P., Ouspenski, I., & Strunnikov, A. (2007). Condensin function at centromere chromatin facilitates proper kinetochore tension and ensures correct mitotic segregation of sister chromatids. Genes to cells: devoted to molecular & cellular mechanisms, 12(9), 1075–1090. https://doi.org/10.1111/j.1365-2443.2007.01109.x

Zakian, V. A., Brewer, B. J., & Fangman, W. L. (1979). Replication of each copy of the yeast 2 micron DNA plasmid occurs during the S phase. Cell, 17(4), 923–934. https://doi.org/10.1016/0092-8674(79)90332-5

